# Urethane Anesthesia Exhibits Neurophysiological Correlates of Unconsciousness and is Distinct from Sleep

**DOI:** 10.1101/2021.09.21.461281

**Authors:** Alejandra Mondino, Joaquín González, Duan Li, Diego Mateos, Lucía Osorio, Matías Cavelli, Alicia Costa, Giancarlo Vanini, George Mashour, Pablo Torterolo

## Abstract

Urethane is a general anesthetic widely used in animal research. It is unique among anesthetics because urethane anesthesia alternates between macroscopically distinct electrographic states: a slow-wave state that resembles NREM sleep (NREMure), and an activated state with features of both REM sleep and wakefulness (REMure). However, the relationship between urethane anesthesia and physiological sleep is still unclear. In this study, electroencephalography (EEG) and electromyography were recorded in chronically prepared rats during natural sleep-wake states and during urethane anesthesia. We subsequently analyzed the EEG signatures associated with the loss of consciousness and found that, in comparison to natural sleep-wake states, the power, coherence, directed connectivity and complexity of brain oscillations are distinct during urethane. We also demonstrate that both urethane states have clear EEG signatures of general anesthesia. Thus, despite superficial similarities that have led others to conclude that urethane is a model of sleep, the electrocortical traits of depressed and activated states during urethane anesthesia differ from physiological sleep states.

## Introduction

Urethane is an anesthetic that is widely used in animal research because of its unique characteristics. It induces long-term narcosis and immobilization with minimal effect on physiological variables (Maggi & Meli, 1986). Furthermore, it produces a spontaneous cycle between two different states: one characterized by slow wave oscillations in the electroencephalogram (EEG), similar to natural non-rapid eye movements (NREM) sleep, and an activated state with rhythmic activity in the slow theta range that qualitatively resembles the EEG pattern either of wakefulness (W) or REM sleep (Détári & Vanderwolf, 1987; Murakami *et al*., 2005; Pagliardini *et al*., 2013; Yagishita *et al*., 2020). Because of the resemblance to a sleep cycle, urethane anesthesia has been proposed as a pharmacological model to study sleep (Horner & Kubin, 1999; Clement *et al*., 2008; Pagliardini *et al*., 2013).

Whether the activated state of urethane anesthesia reflects a wake-like or a REM sleep-like state remains an open question. On the one hand, some authors have proposed that this state corresponds to a reduced level of anesthesia (or a W-like state) because nociceptive stimuli evoke a shift from the NREM-like state to the activated state (Pagliardini *et al*., 2013; Neves *et al*., 2018). Other findings supporting this hypothesis are that rapid eye movements do not occur during the activated state and increasing doses of urethane decrease the relative amount of time spent in that state well as that (Clement *et al*., 2008). In addition, while midline thalamic neuronal firing increases during REM sleep (up to five times higher in comparison to NREM sleep), this increase was less pronounced during the activated urethane state compared to the NREM-like state (Hay, 2021). On the other hand, there is evidence that this activated state resembles REM sleep, including: 1) there are no differences between both urethane states in a withdrawal response to a noxious stimulus (Clement *et al*., 2008); 2) the alternation between the activated and NREM-like state under urethane anesthesia resembles the natural NREM and REM sleep cycle (Clement *et al*., 2008; Pagliardini *et al*., 2013); 3) muscle tone is reduced during the activated state in comparison to the NREM-like state (Clement *et al*., 2008); and 4) there is a similar modulation of breathing between the REM-like state and REM sleep (Pagliardini *et al*., 2012).

In the past years, behavior-independent measures of consciousness and unconsciousness have been developed (Mashour & Hudetz, 2018), providing a novel opportunity to study urethane states. For instance, neural oscillations differ between conscious and unconscious states. Specifically, gamma oscillations (30-150 Hz), hypothesized to relate to cognitive processes, appear during W and REM and are reduced during NREM or anesthesia (Hudetz *et al*., 2011; Cavelli *et al*., 2015; Mondino *et al*., 2020). Conversely, delta oscillations (0.1-4 Hz) occur both during NREM and anesthesia, possibly disrupting the neural interactions supporting consciousness (Steriade *et al*., 1993; Pigorini *et al*., 2015; Tononi *et al*., 2016; Arena *et al*., 2021). In addition, synchronization between high frequency neural oscillations at different brain areas is another major correlate of consciousness (Varela *et al*., 2001; Melloni *et al*., 2007). Gamma oscillations are synchronous during alert and awake states, but they decouple during NREM, anesthesia and even REM sleep (Lee *et al*., 2013; Cavelli *et al*., 2015; Pal *et al*., 2016; Castro-Zaballa *et al*., 2018; Mondino *et al*., 2020). The directionality of the connectivity is another important feature to consider, as feedforward and feedback interactions between anterior and posterior cortices decrease during sleep and different anesthetics (Pal *et al*., 2016). Finally, the complexity of brain oscillations changes between conscious and unconscious states, suggesting that complex neuronal interactions are necessary for awareness (Schartner *et al*., 2015; Mateos *et al*., 2017; Schartner *et al*., 2017; Demertzi *et al*., 2019; Gonzalez *et al*., 2019; Gonzalez *et al*., 2020; Sarasso *et al*., 2021).

In the present report, we characterize the EEG correlates of urethane anesthesia. Specifically, we compared electrocortical activity (power, coherence, directionality and complexity) between physiologic sleep-wake states and NREM-like urethane (NREMure) and REM-like urethane (REMure) states of anesthesia. We found that the urethane states show a consistent decline in the EEG correlates of consciousness, suggesting that—despite the cyclic alternation between EEG profiles—urethane induces sustained unconsciousness. In addition, we demonstrate well-defined EEG differences between NREMure and REMure and their correspondent sleep state. Based on these differences we conclude that, despite superficial similarities, urethane anesthesia is not an accurate model of sleep.

## Materials and Methods

### Experimental animals

Nine Wistar male adult rats (270–300 g) were studied. Veterinarians of the institution certified that the rats were in good health at the time of the study. All experimental procedures were conducted in agreement with the National Animal Care Law (#18611) and with the “Guide for the care and use of laboratory animals” (8th edition, National Academy Press, Washington, DC, USA, 2010). Furthermore, the Institutional Animal Care Committee approved the experimental procedures (No 070153-000332-16). Adequate procedures were taken to minimize animals’ pain, discomfort, and stress.

We utilized the minimum number of animals necessary to acquire reliable scientific information. The rats were housed in groups of four to five per cage prior to the experiments. Before and during the experiments they were kept in a temperature-controlled (21–24 °C) room, on a 12:12 h light/dark cycle (lights on at 6.00 a.m.) with food and water available *ad libitum*.

### Surgical Procedures

We employed the same surgical procedure that we used in our previous studies (Gonzalez *et al*., 2018; Mondino *et al*., 2019; Mondino *et al*., 2020). Anesthesia was induced with a combination of ketamine and xylazine (90 mg/kg; 5 mg/kg, i.p., respectively). Ketoprofen, 1 mg/kg subcutaneous was administered to provide analgesia. Rats were positioned in a stereotaxic frame; the skull was exposed, and seven stainless-steel screw electrodes were implanted in the skull in order to record the intracranial EEG or electrocorticogram. The electrodes were localized above the right and left primary motor cortex (M1), primary somatosensory cortex (S1) and secondary visual cortex (V2). Another electrode was placed in the right olfactory bulb (OB). Finally, a reference electrode was implanted in the cerebellum. The electromyogram (EMG) was recorded by means of a bipolar electrode inserted into the neck muscles. A representation of the electrode positions is shown in Figure 1A. The electrodes were soldered to a socket and fixed to the skull with dental cement. A topical antibiotic was applied to the margins of the incision. Animals were allowed to recover from the surgical procedure for seven days. Thereafter, they were housed independently in transparent cages (40 × 30 × 20 cm) and habituated for five days to the recording condition in a sound-attenuated chamber.

**Figure 1.**
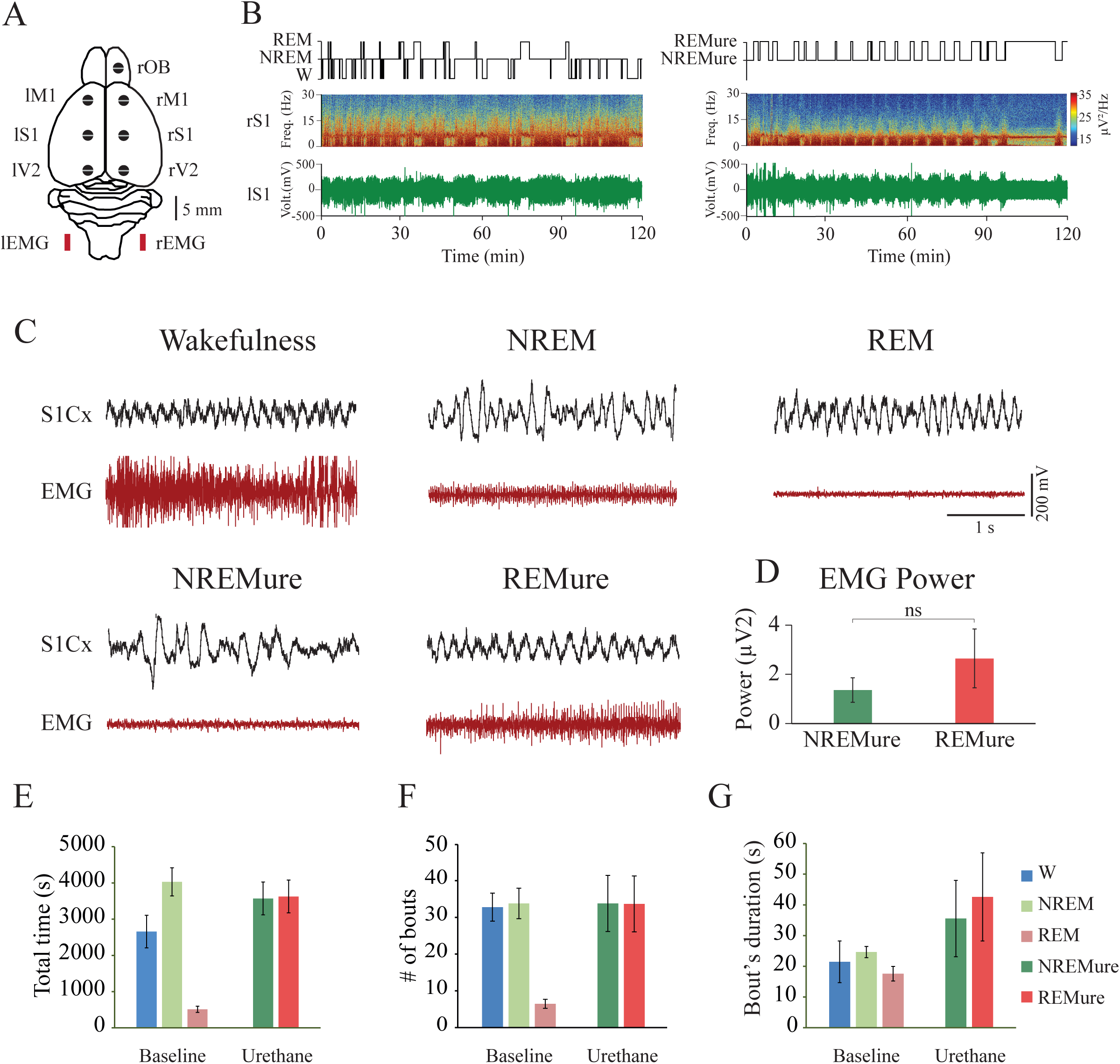
Sleep–wake and urethane states in the rat. (**A**) Schematic representation of the electrode position in the brain of the rat. OB, olfactory bulb; M1, primary motor cortex; S1, primary somato-sensory cortex; V2, secondary visual cortex; r, right; l, left. (**B**) Hypnogram (top) according to visually scored behavioral states and spectrogram (0.1 to 30 Hz) of one representative animal. Left panel shows baseline recordings and right panel shows recordings under urethane anesthesia. During W, REM sleep and REMure theta activity (4–10 Hz) in the spectrogram can be readily observed. **(C)** Representative electroencephalogram (EEG) of the somatosensory cortex (S1Cx) and neck electromyogram (EMG) recordings during W, NREM sleep, REM sleep, NREMure and REMure. Similarities between the sleep and anesthetized states can be observed with delta (0.5 – 4 Hz) oscillations characterizing NREM sleep and NREMure and theta (4-10 Hz) and high frequency oscillations characterizing W, REM sleep and REMure. Notice that REMure doesn’t show the REM sleep characteristic atonia. **(D)** Electromyography power during NREMure and REMure, both states showed no significant (ns) differences. **(E)** Mean and SEM of the total time spent in each behavioral state during baseline recordings (left) and urethane anesthesia (right). **(F)** Mean and SEM of the number of bouts of each behavioral state during baseline recordings (left) and urethane anesthesia (right). **(G)** Mean and SEM of the duration of bouts of each behavioral state during baseline (left) and urethane anesthesia (right).

### Sleep Recordings

Experimental sessions were performed between 9:00 AM and 1:00 PM. The recordings were carried out using a rotating connector that allows the rats to freely move within the recording chamber. Bioelectric signals were amplified (×1000), band-pass filtered (0.1–500 Hz), sampled (1024 Hz, 16 bits) and acquired using using DASYlab, (National Instruments, Austin, TX, US).

Baseline polysomnographic recordings were performed for two hours. Thereafter, the rats were anesthetized with urethane (1.2 to 1.5 g/kg, i.p.), and recorded for another two hours. During anesthesia, animals were warmed with a heating pad at 37°C. Video recordings were also performed during the recording sessions.

States of W, NREM and REM sleep were manually scored using Spike 2 software version 9.04 (Cambridge Electronic Design, Cambridge, UK) in 5s epochs. W was defined by low-amplitude, high-frequency EEG activity accompanied by high EMG activity. NREM sleep was defined by high-amplitude slow waves as well as sleep spindles associated with a reduced muscular tone, while REM sleep was identified by low-amplitude, fast waves with a regular theta rhythm, particularly in the posterior cortex, and by an EMG atonia except for occasional twitches. During urethane anesthesia two states were defined according to the EEG profile. NREMure displayed high-amplitude low-frequency EEG similar to NREM sleep, whereas REMure was characterized by low-amplitude and high-frequency EEG associated with regular theta rhythm. These states were also scored in 5s epochs. We determined the total time spent in each state as well as the number and duration of epochs.

### EEG signal analysis

We analyzed all EEG epochs without artifacts or state transitions. The frequency bands were defined as delta (1-4 Hz), theta (4-10 Hz), sigma (10-15 Hz), Beta (15-30 Hz), low gamma (LG, 30-45 Hz), high gamma (HG, 55-95 Hz) and high frequency oscillations (HFO, 105-145, and 155-195 Hz). Frequencies around the alternating current (50 Hz in Uruguay) and harmonics were excluded from the analysis.

The power spectrum was calculated by means of the *pwelch* built-in function on Matlab (version 2020a; MathWorks, Inc., Natick, MA) with a 5s window size, half-window overlap, frequency sample of 1024 Hz and a resolution of 0.5 Hz. Absolute power values were normalized for each animal as the mean power of each frequency band divided by the total power (i.e., sum of the power of each frequency band) (Mondino *et al*., 2020; González *et al*., 2021). Spectral coherence was determined as a measure of undirected functional connectivity between EEG electrodes (Bullock & McClune, 1989), using the same procedure as Mondino et al. (2020). We determined the coherence between interhemispheric and intrahemispheric adjacent electrodes (the distance between neocortical contiguous electrodes was 5 mm). The *mscohere* built-in Matlab function was employed for this analysis with a 5s window size, half-window overlap, frequency sample of 1024 Hz and a resolution of 0.5 Hz. Coherence values were normalized by the Fisher’s z-transform to get the z’coherence. For the power and coherence analysis, the rOB, rV2, lM1 and lS1 had to be excluded in one rat during the baseline recordings, and rV2 and lM1 during anesthetized recordings due to artifacts that made processing unfeasible. Additionally, rS1-rV2 z’coherence of another rat was excluded from the analysis because it was a clear outlier with extremely high values for all the frequency bands.

Normalized symbolic transfer entropy (NSTE) was used to determine directed connectivity between the rM1 (anterior) and rV2 (posterior) cortices (Borjigin *et al*., 2013; Lee *et al*., 2013; Pal *et al*., 2016). rM1 and rV2 EEG signals were filtered into the frequency bands described above and segmented into non-overlapped 5s windows. We used the same strategy for parameter selection as in our previous studies (Li *et al*., 2017; Mondino *et al*., 2021). The embedding dimension was fixed at d*E* = 3 and we used the time delay (τ) = 128, 51, 34, 17, 11, 5, 3 corresponding respectively to delta, theta, sigma, beta, LG, HG and HFO. In every window, we examined the transfer time (δ) = 1-100 (corresponding to 1-100 ms approximately) and chose the one that generated the maximum feedback (anterior to posterior) and feedforward (posterior to anterior) NSTE, respectively.

Finally, we evaluated the complexity of the EEG signal by means of Lempel-Ziv Complexity (LZC) (Lempel & Ziv, 1976). This is an information theoretic measure based on Kolmogorov complexity, that calculates the minimal “information” contained in the sequence (Cover & Thomas, 2006). LZC has been shown to be an important tool to explore the temporal complexity of brain activity (Schartner *et al*., 2015; Hudetz *et al*., 2016). To estimate the complexity of a time series 𝒳*(t)* ≡ {*x*_*t*_; *t* =1, …, *T*}, a sequence *X(t)* is parsed into a number of words *W* by considering any subsequent series that has not yet been encountered as a new word. The LZC *c*_*LZ*_ is the minimum number of *W* required to reconstruct the information contained in the original time series. Continuous sequences such as an EEG signal, need to be discretized to analyze LZC. Different methods can be employed for this purpose and could lead to slightly differing results. In this work, we employed the binarization by mean-value (referred to as bin LZC) to calculate the complexity of each EEG channel. Although Lempel and Ziv originally developed the method for binary sequences, this approach can be easily generalized to multivariate discrete processes by extending the alphabet size (Zozor *et al*., 2005). Consider an *m*-dimensional stationary process *X*(*m*), that generates the sequences *x*_*t,i*_ = *x*_1,*i*_ ,…, *x*_*T,i*_ with *i* = 1,…,*m* each one of them from an alphabet of *α* symbols. Let *z*_*t*_ = z_1_ ,… ,*z*_*T*_ to be a new sequence defined over an extended alphabet of size *α*_*m*_

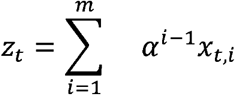

Using this approach, we also calculated the global complexity of neurophysiologic activity, referred as joint Lempel–Ziv complexity (JLZC) of the *x*_*i*_-sequence as the complexity of the new sequences *z*_*t*_ *C*_*LZ*_(*x*_*t,i*_) = *C*_*LZ*_(*z*_*t*_). In this case, *x*_*i,t*_ is the EEG signal per channel and *z*_*t*_ is a new sequence that represents the information from all channels. The JLZC was binarized (*α* = 2) using the mean value (Zozor *et al*., 2005).

### Statistical analyses

Statistical analyses were done with PRISM v9.0 (GraphPad Software Inc., La Jolla, CA). All data were presented as mean ± standard error of the mean (SEM). Comparisons between W, NREMure and REMure were performed by means of a repeated measures one-way ANOVA and Holm-Sidak as a *post hoc* test; adjusted p value < 0.05 was considered significant. Two-tailed paired Student t tests were used to determine differences between NREM and NREMure as well as REM and REMure. p values were also corrected for multiple comparisons using Holm-Sidak test.

## Results

### Rats under urethane anesthesia cycle between two different states

A representative hypnogram, spectrogram (power spectrum as a function of time), as well as the EEG recordings of baseline and following urethane anesthesia during two hours of recordings are shown in Figure 1B. During baseline, rats exhibit the states of W, NREM and REM sleep, while under urethane anesthesia they cycled between two different states, a slow wave activity state (NREMure), and an activated state (REMure). In Figure 1C, the distinctive characteristics of the EEG and EMG of each state can be observed. Although muscle activity in most of the rats was higher in REMure than during NREMure, no difference in the EMG power was observed when the whole population was analyzed (Figure 1D). As can be appreciated in Figure 1E-G during the baseline recordings rats spent 2702 ± 422.6 s (37.52% of the total time) in W, 3977 ± 367.8 s (55.23%) in NREM sleep and 522 ± 76.52 s (7.25%) in REM sleep. If we consider the total amount of sleep, NREM sleep represented 88.4% while REM sleep represented 11.6%. In average, the number of bouts of W was 32.78 ± 3.79 with a mean duration of 21.45 ± 7.08 s, the number of NREM bouts was 33.78 ± 4.15 with a duration of 24.65 ± 1.82 s, and the number of REM bouts was 6.44 ± 1.23 and lasted 17.58 ± 2.36 s. After urethane injection, rats spent a similar amount of time in NREMure and REMure; 3292 ± 434.15 s (45.73% of total time) and 3908 ± 434.15 s (54.27%), respectively [t (8) = 0.708; p = 0.4985]. The number of bouts of NREMure was 33.78 ± 7.63, and of REMure 33.67 ± 7.59 [t (8) = 0.00; p >0.9999]. There was no difference in bout duration between the two states (35.54 ± 37.32 s for NREMure and 42.60 ± 14.36 for REMure [t (8) = 1.067; p = 0.3172]).

### Urethane increases low and decreases high frequency bands of the EEG

To study brain oscillations, we analyzed the EEG power spectrum from OB, M1, S1 and V2. We compared both urethane states to the physiological W to identify EEG features of unconsciousness during urethane (Figure 2A). Detailed statistics are shown in Supplementary Table 1. We found that delta oscillations increased in all cortical locations during NREMure compared to either REMure or W. In addition, W had more power in the rest of the bands compared to NREMure. This can be fully appreciated looking at low-gamma, high-gamma and HFO bands, which decrease in all cortical locations during NREMure. In contrast, the distinction between W and REMure was less marked, and was evidenced by a power decrease in the frequencies above low-gamma. In particular, HFOs decreased during REMure in all cortical locations (significant in OB, M1 and S1), while low and high gamma also significantly decreased in M1 and S1. We also observed a noticeable decrease in the theta peak frequency during REMure compared to W and REM [Mean ± SEM, 4.75 ± 0.16 for REMure, 6.87 ± 0.12 for REM sleep, and 6.94 ± 0.15 for W; [F (2,7) = 79.40, p <0.0001]. For this analysis, we calculated the differences between the maximum value in each state in V2, where theta oscillations are fully expressed.

**Figure 2.**
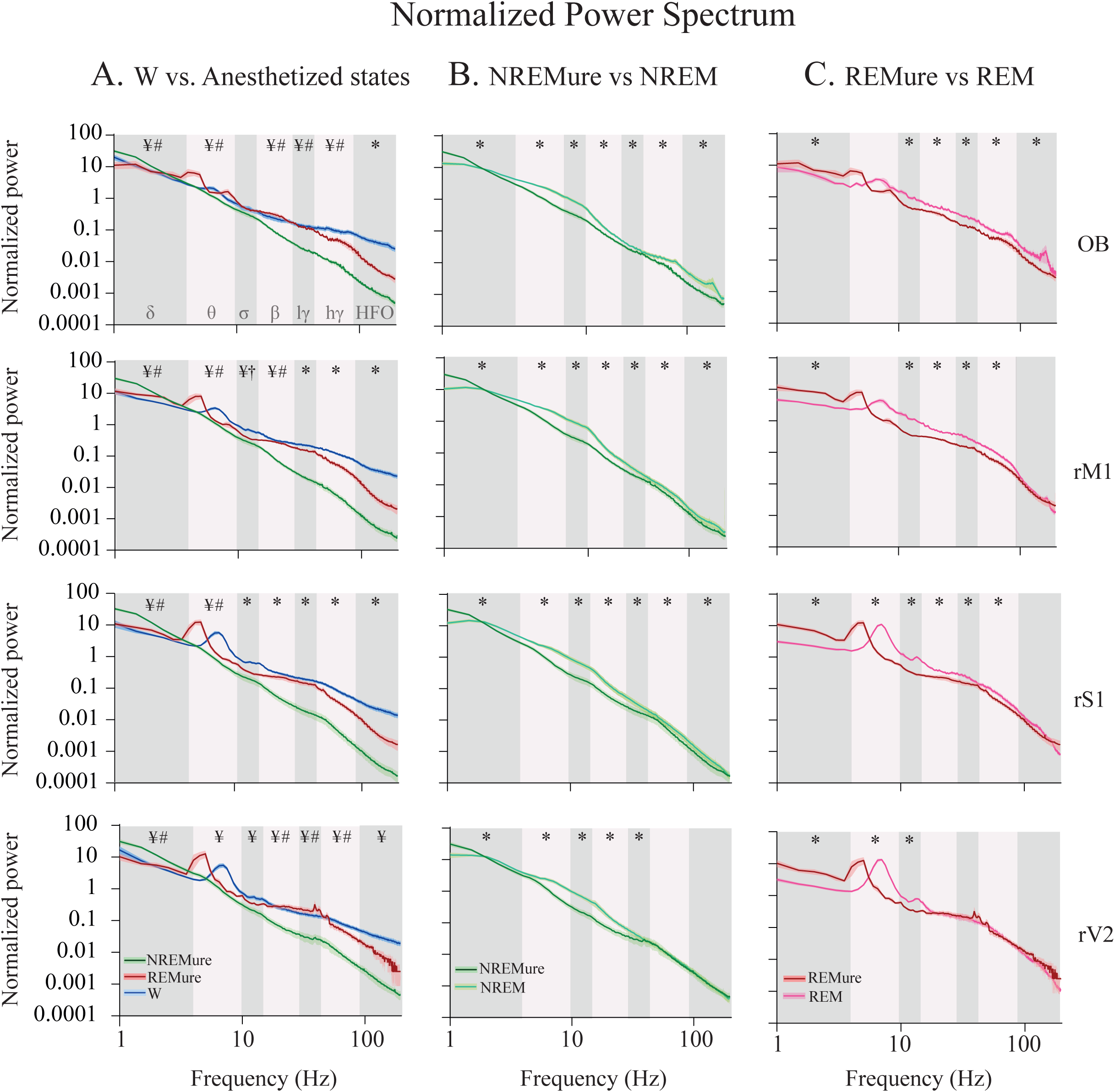
Power spectral profiles. **(A)** Mean and SEM (shadow) normalized power spectral profiles of the right hemisphere during wakefulness (W) and the anesthetized states (NREMure and REMure). Significant diffrerences are indicated by symbols: ¥, W Vs. NREMure; †, W Vs. REMure; #, NREMure Vs. REMure; *, differences among all conditions. **(B)** Mean and SEM (shadow) of normalized power spectral profiles of the right hemisphere during NREM sleep and NREMure. *, indicates significant differences. **(C)** Mean and SEM (shadow) normalized power spectral profiles of the right hemisphere during REM sleep and REMure. *, indicates significant differences. The frequency bands used for the statistical analysis are indicated by colors in the background of the chart. OB, olfactory bulb; M1, primary motor cortex; S1, primary somato-sensory cortex; V2, secondary visual cortex; W, wakefulness; r, right; l, left; l; δ, delta; θ, theta; σ, sigma; β, beta; lγ, low gamma or LG; hγ, high gamma or HG; HFO, high frequency oscillations.

Next, we compared both urethane states to study how the various brain oscillations differ between them; statistics are shown in Supplementary Tables 1 and 2. Importantly, we found that delta power was higher during NREMure in all locations. Similar to W, REMure showed higher power in all frequency bands above delta, which was significant for theta in OB, M1, S1; for sigma in S1; and for Beta, LG, HG and HFO in all locations except for HFO in the visual cortex.

To complete the characterization of brain oscillations under urethane anesthesia, we compared each urethane state to its physiological analog; i.e, NREMure vs NREM, and REMure vs REM. Statistics are shown in Supplementary Table 3. Comparing each pair, we found that delta oscillations are larger under both urethane states compared to their physiological counterparts in all cortical sites. The rest of the frequency bands showed a tendency to decrease under urethane. For instance, all bands (>delta) were reduced in NREMure compared to NREM (except for HG and HFO in V2), the same occurred comparing REMure to REM; the exception was HFO in the neocortex as well as beta and gamma bands in V2.

In summary, urethane promotes slow delta oscillations, while reducing higher frequency bands, especially gamma and HFOs.

### Urethane alters corticocortical synchronization

For a more complete characterization of urethane’s neurophysiological correlates, we analyzed the spatial synchronization between neural oscillations. We first analyzed the interhemispheric z’coherence comparing between W and the urethane states (Figure 3A). The complete statistical analysis is shown in Supplementary Table 4A and 5. We found that during NREMure, the inter-hemispheric delta coherence increased with respect to the other states, which was significant in rM1-lM1 comparing NREMure to REMure or W, and rS1-lS1 comparing NREMure to W. Moreover, higher frequency bands were more coherent during W than during both urethane states. For instance, rS1-lS1 theta and sigma coherences decreased during REMure compared to W, while rS1-lS1 HG and HFO decreased for both urethane states compared to W.

**Figure 3.**
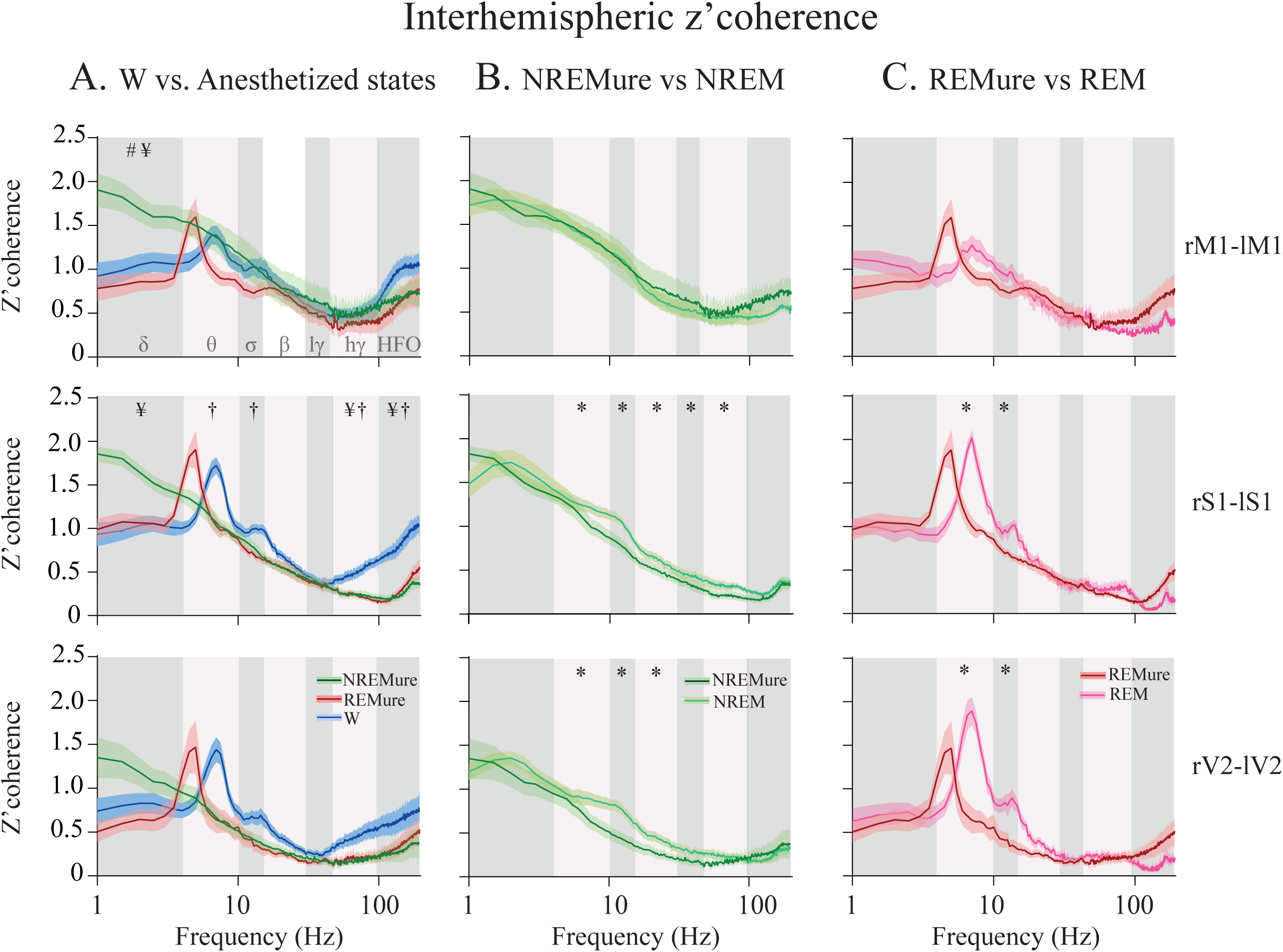
Interhemispheric z’coherence. **(A)** Mean and SEM (shadow) interhemispheric z’coherence during wakefulness (W) and the anesthetized states (NREMure and REMure). Significant diffrerences are indicated by symbols: ¥, W Vs. NREMure; †, W Vs. REMure; #, NREMure Vs. REMure; *, differences among all conditions. **(B)** Mean and SEM (shadow) interhemispheric z’coherence during NREM sleep and NREMure. *, indicates significant differences. **(C)** Mean and SEM (shadow) interhemispheric z’coherence during REM sleep and REMure. *, indicates significant differences. M1, primary motor cortex; S1, primary somato-sensory cortex; V2, secondary visual cortex; W, wakefulness; r, right; l, left; l, δ, delta; θ, theta; σ, sigma; β, beta; lγ, low gamma or LG; hγ, high gamma or HG; HFO, high frequency oscillations

Comparing each urethane state to its natural analog, we found differences suggesting that inter-hemispheric synchronization is compromised compared to physiological sleep states. Details of these statistics are shown in Supplementary Table 7. Comparing NREMure to NREM, we found that theta, sigma and beta coherences decreased in both rS1-lS1 and rV2-lV2, and that LG and HG coherences decreased in rS1-lS1 (Figure 3B). The REMure vs REM comparison showed that theta and sigma oscillations are also less coherent during REMure in rS1-lS1 and rV2-lV2 (Figure 3C).

To more fully characterize how urethane alters cortical synchronization, we also studied the intra-hemispheric z’coherence between adjacent electrodes on the right hemisphere. We found that delta coherence increased for both urethane states in OB-M1 compared to W. Moreover, HG coherence between M1-S1 and S1-V2 decreased during NREMure compared to W, while HFO S1-V2 coherence also decreased during both urethane states compared to W (Figure 4A). When we compared each urethane state to its analog, we found no significant differences (Figure 4B and C).

**Figure 4.**
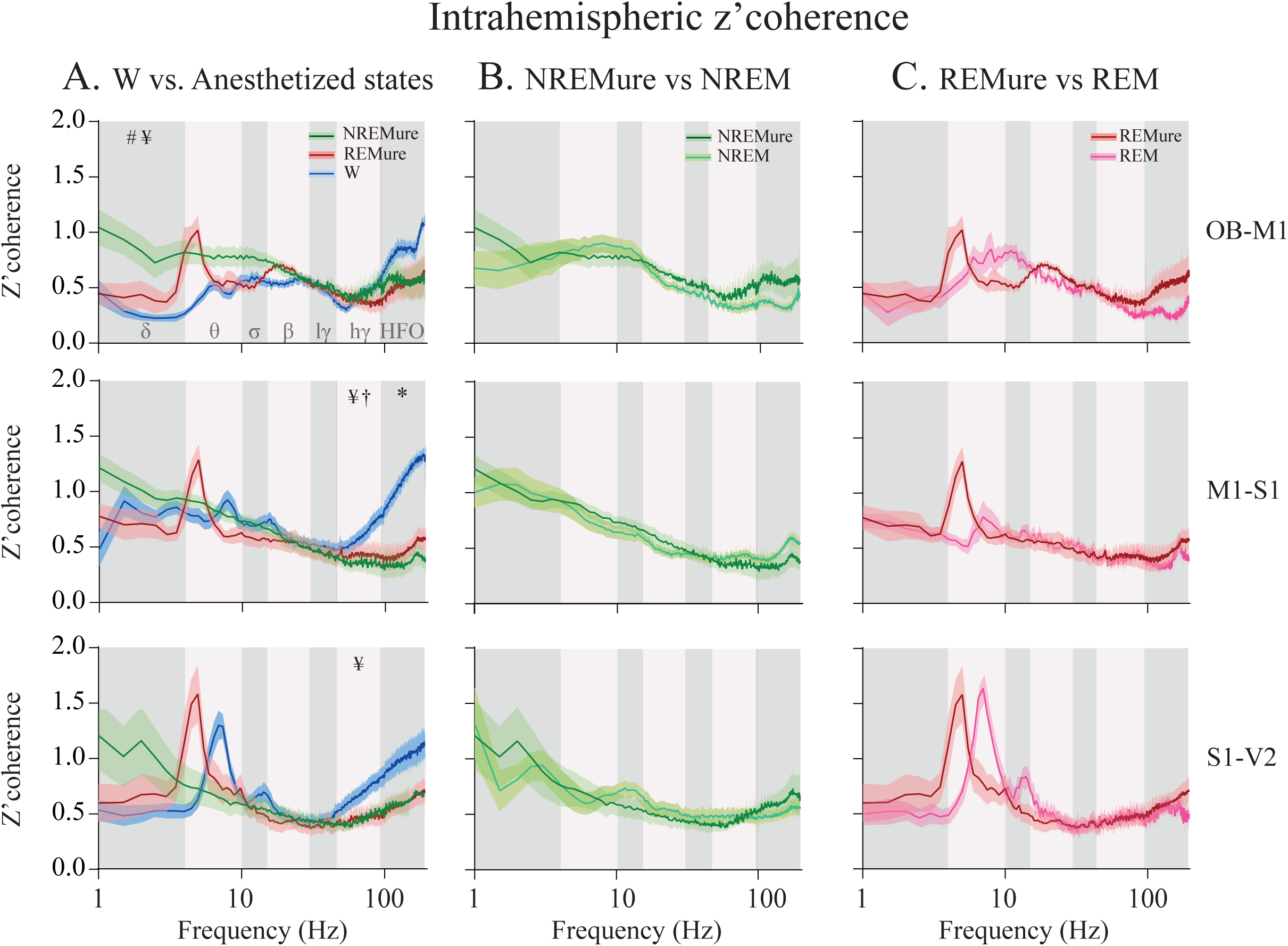
Right Intrahemispheric z’coherence. **(A)** Mean and SEM (shadow) right intrahemispheric z’coherence during wakefulness (W) and the anesthetized states (NREMure and REMure). Significant diffrerences are indicated by symbols: ¥, W Vs. NREMure; †, W Vs. REMure; #, NREMure Vs. REMure; *, differences among all conditions. **(B)** Mean and SEM (shadow) right intrahemispheric Z’coherence during NREM sleep and NREMure. *, indicates significant differences. **(C)** Mean and SEM (shadow) right intrahemispheric Z’coherence during REM sleep and REMure. OB, olfactory bulb; M1, primary motor cortex; S1, primary somato-sensory cortex; V2, secondary visual cortex; W, wakefulness; r, right; l, left; l, δ, delta; θ, theta; σ, sigma; β, beta; lγ, low gamma or LG; hγ, high gamma or HG; HFO, high frequency oscillations

Thus, these results show that urethane promotes synchronized delta waves between areas, while higher frequency synchronization, such as gamma, decreases during both urethane states.

### Urethane compromises feedback connectivity

To evaluate the directed connectivity between the rM1 (anterior) and rV2 (posterior) cortices, we calculated feedback and feedforward NSTE for each frequency band. As is illustrated in Figure 5, when comparing W to the anesthetized states, both feedback and feedforward NSTE were affected by behavioral states in the theta frequency band [F(2,8) = 9.523, p = 0.0081 and F(2,8) = 7.433, p = 0.017 respectively]. NREMure had a lower connectivity than W and REMure in both feedback (p = 0.049 and 0.026, respectively) and feedforward directions (p = 0.029 and 0.002, respectively). No differences were found between W and REMure for feedback or feedforward connectivity (p = 0.140 and 0.499 respectively). Additionally, feedback but not feedforward connectivity was affected by the states in HG frequency band [F(2,8) = 6.620, p = 0.0136 and F(2,8) = 0.783, p = 0.425, respectively]. W feedback NSTE was higher than REMure (p = 0.049) but not than NREMure (p = 0.140). Lastly, both anesthetized states did not differ in HG feedback NSTE (p = 0.361). Delta, sigma, beta, LG and HFO frequency bands did not show differences between W and the anesthetized states neither for the feedback NSTE [F(2,8) = 4.536, p = 0.055, F(2,8) = 2.436, p = 0.150, F(2,8) = 1.056, p = 0.348, F(2,8) = 0.344, p = 0.621 and F(2,8) = 3.522, p = 0.078 respectively] nor for the feedforward NSTE [F(2,8) = 0.077, p = 0.827, F(2,8) = 4.479, p = 0.063, F(2,8) = 0.601, p = 0.487, F(2,8) = 0.739, p = 0.459 and F(2,8) = 3.509, p = 0.077 respectively].

**Figure 5.**
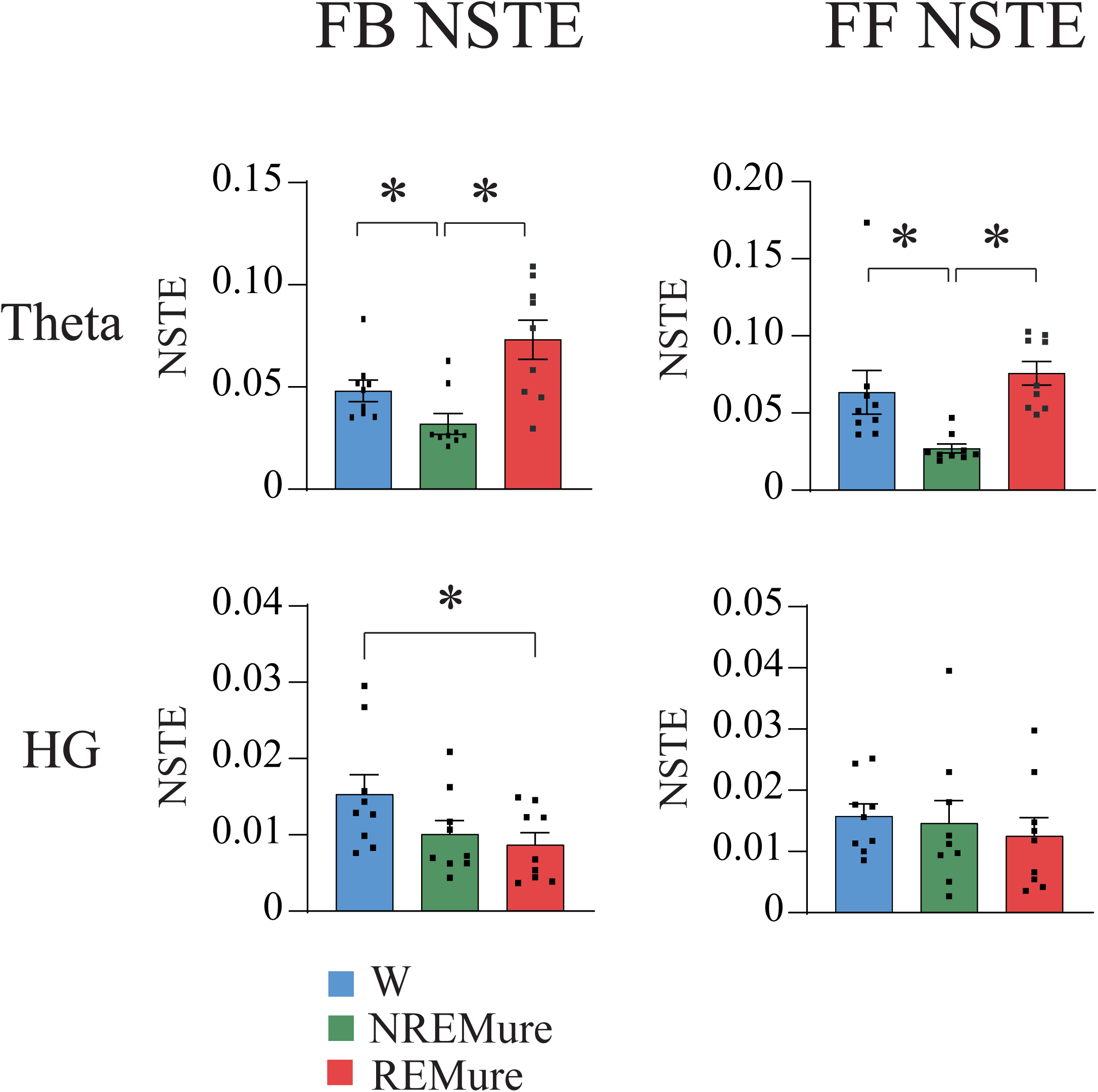
Normalized Transfer Entropy. Feedback (FB) and Feedforward (FF) Normalized transfer entropy (NSTE) for Theta and HG frequency bands during W and the anesthetized states. Frequency bands with no significant differences between states are not shown. *, indicates significant differences. Comparisons of NSTE between sleep states and its analogous during anesthesia can be find in Supplementary Table 9.

When comparing the anesthetized states with its analogous state during sleep, we did not see any differences in feedback or feedforward connectivity. The statistics are shown in Supplementary Table 9.

### Urethane decreases EEG complexity

We studied the complexity of the EEG signal during each state by means of the binarized Lempel Ziv Complexity (LZC). First, we analyzed the complexity of the whole signal (1-195 Hz). In all the electrode locations the ANOVA showed significant differences between W and urethane states [rOB. F (2,7) = 25.48 p = 0.0002], [rM1 F (2,8) = 46.53 p <0.0001], [rS1 F (2,8) = 69.89 p <0.0001] and [rV2 F (2,8) = 64.02 p = <0.0001]. As can be observed in Figure 6A, in rOB, rM1, rS1 and rV2 the complexity of the EEG signal during NREMure was lower than during W (p = 0.0022, <0.0001 and <0.0001, respectively) and REMure (p = 0.0036, 0.0004, <0.0001 and <0.0001, respectively). Additionally, W had a higher complexity than REMure in rOB, rM1 and rS1 (p = 0.0064, 0.0139 and 0.0042, respectively) but not in rV2 (p = 0.0849). When comparing the NREMure with NREM sleep, no differences in complexity were found in any of the electrode locations (rOB p = 0.1038, rM1 p = 0.0691, rS1 p = 0.6627, rV2 p = 0.0835). The same occurred between REMure and REM sleep (rOB p = 0.4280, rM1 p = 0.5359, rS1 p =0.2824, rV2 p = 0.3638). Because sleep is characterized by slow oscillations, we measured the LZC for the signal between 1 to 15 Hz. Like the whole signal analysis, the ANOVA showed significant differences between W and the anesthetized states [rOB F (2,7) = 13.75 p = 0.0007], [rM1 F (2,8) = 43.30 p = <0.0001], [rS1 F (2,8) = 94.97, p <0.0001] and [rV2 F (2,8) = 71.90, p <0.0001]. In all cortical areas NREMure had a lower complexity than W (rOB p = 0.0027, rM1 p <0.0001, rS1 p <0.0001, rV2 p <0.0001) and REMure (rOB p = 0.0144, rM1 p = 0.0036, rS1 p = 0.0005 and rV2 p = 0.0007). In addition, W had a higher complexity than REMure in rM1 (p = 0.0004), rS1 (p <0.0001) and rV2 (p = 0.0001) but not in rOB (p = 0.5278). Additionally, we compared the complexity during the anesthetized states and in analogous states during sleep, observing clear differences between them. For instance, NREMure had lower complexity in the low frequency oscillations than NREM sleep in all electrode locations (rOB p = 0.0007, rM1 p <0.0001, rS1 p <0.0001 and rV2 p <0.0001). Similarly, REMure had lower complexity than REM sleep (rOB p = 0.0008, rM1 p <0.0001, rS1 p <0.0001 and rV2 p <0.0001).

**Figure 6.**
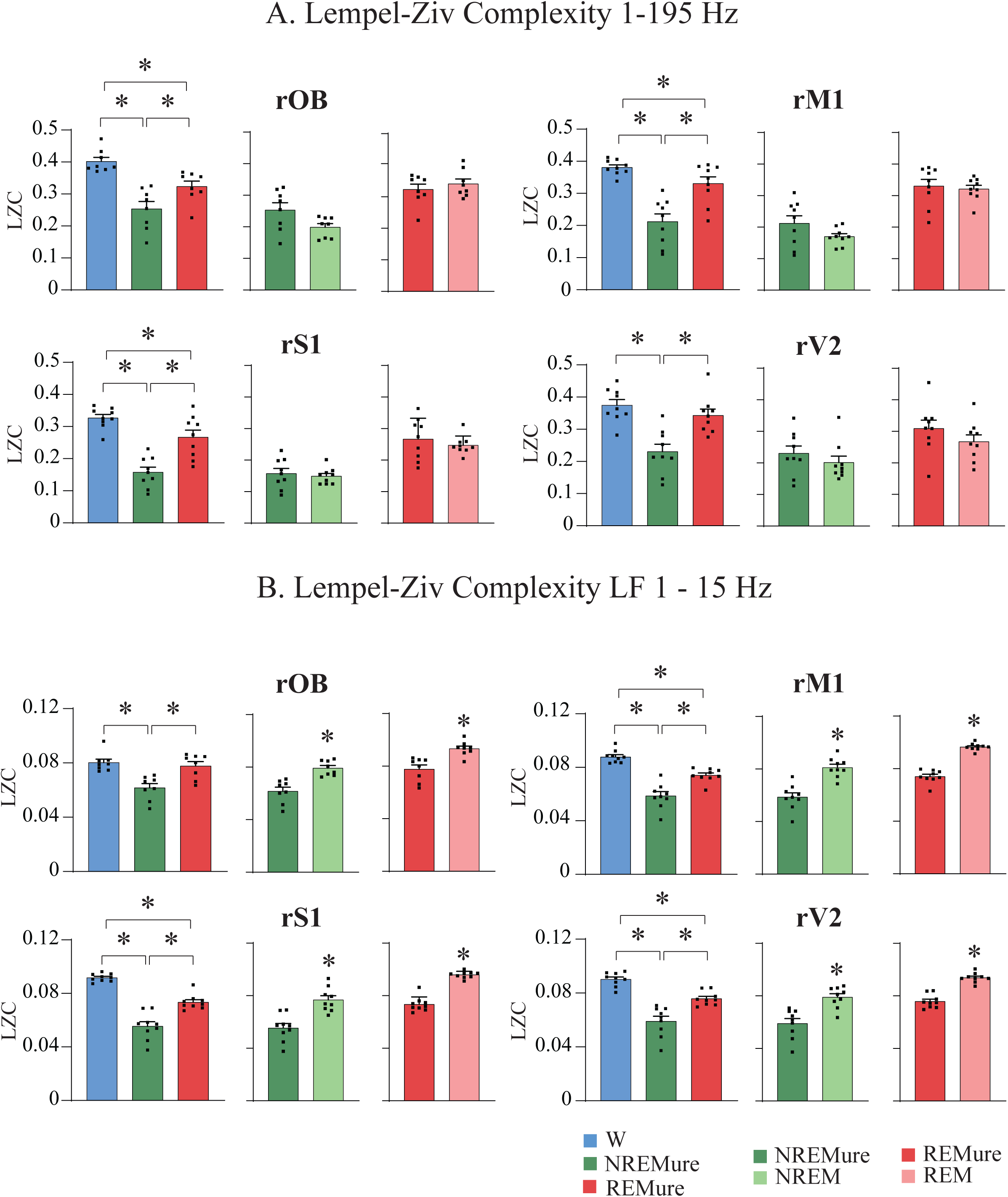
Lempel Ziv Complexity (LZC). **(A)** Lempel Ziv Complexity for frequencies between 1-195 Hz during W and the anesthetized states (Left), NREM sleep and NREMure (middle) and REM sleep and REMure (right) for each electrode localization in the right hemisphere. *, indicates significant differences. **(B)** Lempel Ziv Complexity for frequencies between 1 -15 Hz (low frequencies, LF) during W and the anesthetized states (Left), NREM sleep and NREMure (middle) and REM sleep and REMure (right) for each electrode localization in the right hemisphere. *, indicates significant differences. OB, olfactory bulb; M1, primary motor cortex; S1, primary somato-sensory cortex; V2, secondary visual cortex; W, wakefulness; r, right; l, left.

Thus, unlike directed connectivity measures, complexity analysis distinguished between physiological sleep and urethane-induced states.

### Urethane disrupts whole-brain complexity

Finally, to understand how the complexity of the EEG signal within the whole brain varies between states, we computed a joint Lempel Ziv for 1–195 Hz and for 1 – 15 Hz. The results are displayed in Figure 7. ANOVA demonstrated a clear difference between the complexity during W and the anesthetized states for 1 – 195 Hz F(2,8) = 82.63 p < 0.0001. W had a higher complexity than NREMure p < 0.0001and REMure p = 0.0099. Also, REMure had a higher complexity than NREMure (p = < 0.0001). Similar results were found for the 1 – 15 Hz oscillations F(2,8) = 55.27 p < 0.0001. W had a higher complexity than NREMure p < 0.0001 and REMure p = 0.0099. Furthermore, REMure had a higher complexity than NREMure (p < 0.0009).

**Figure 7.**
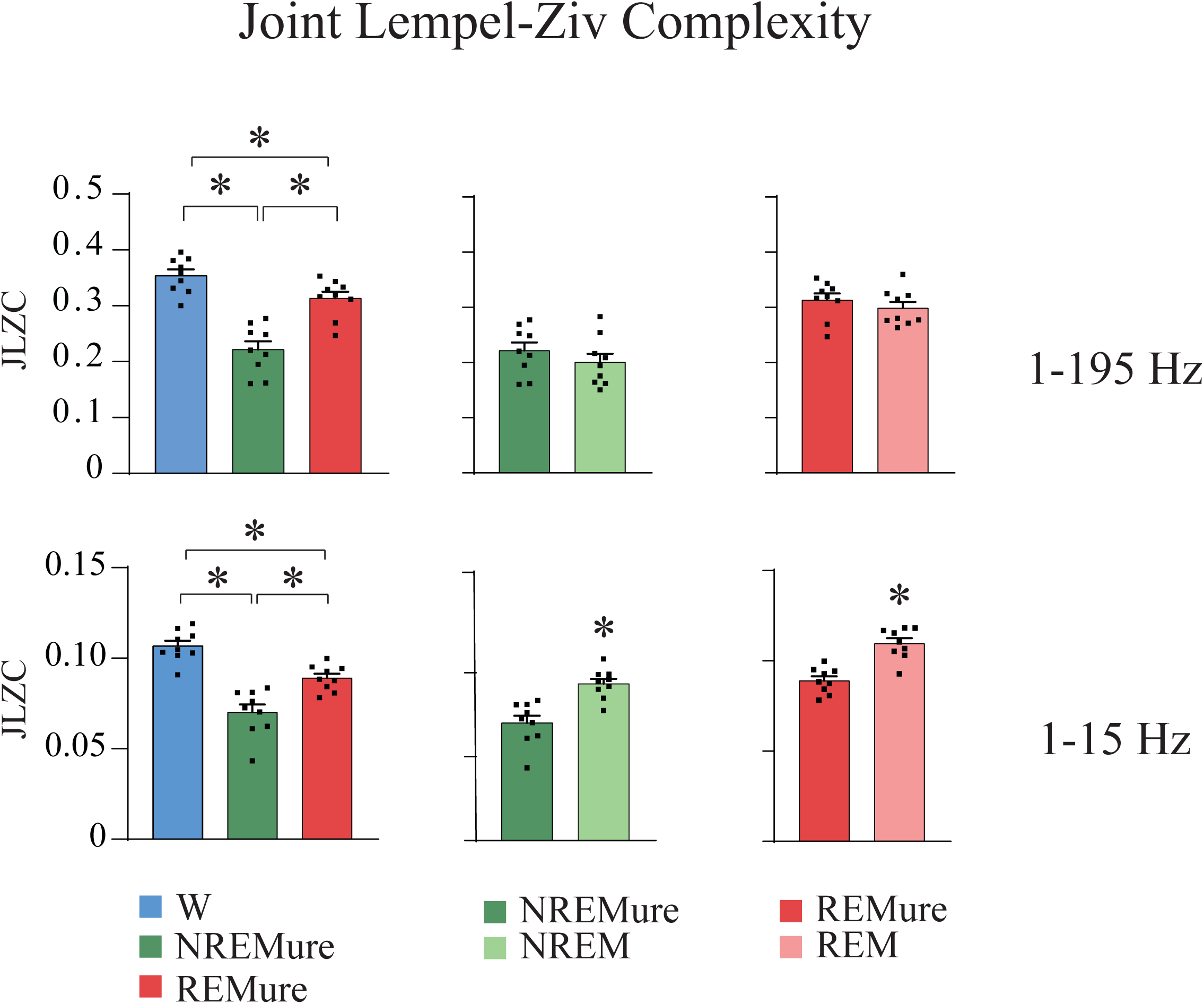
Joint Lempel Ziv Complexity (JLZC). **(A)** Comparison of Lempel Ziv Complexity for frequencies between 1-195 Hz (top) and 1-15 Hz (bottom) for W and the anesthetized states (Left), NREM sleep and NREMure (middle) and REM sleep and REMure (right). *indicates significant differences. OB, olfactory bulb; M1, primary motor cortex; S1, primary somato-sensory cortex; V2, secondary visual cortex; W, wakefulness; r, right; l, left.

Differences between sleep states and anesthetized states were seen for the 1 - 15 Hz analysis NREM sleep had higher complexity than NREMure (p = 0.0002), and REM sleep had higher complexity than REMure (p < 0.0001). For the 1 – 195 Hz analysis these differences were lost (NREM sleep vs NREMure p = 0.2125, and REM sleep vs REMure p = 0.1693).

## Discussion

We performed an extensive analysis of the EEG signal during urethane anesthesia. We demonstrated that, in comparison to W, the EEG signal quantified by different metrics is significantly modified during both urethane-induced states, i.e., NREMure and REMure. In addition, while urethane anesthesia resembles natural sleep, the EEG profiles under urethane anesthesia and sleep are not the same. This brings into question whether urethane can serve as a pharmacological model for sleep.

### General features of urethane-induced states

Urethane evokes a cycle between two states that resemble natural sleep states. However, despite the shared phenotype, our EEG analysis suggests that the underlying oscillations and neurobiology of sleep and urethane anesthesia are different. REMure did not show the characteristic muscle atonia that occurs in REM sleep. In fact, no differences were observed between the two anesthetized states in the total power of the EMG (Figure 1D). This finding is in disagreement with the finding of Clement *et al*. (2008), that observed a decreased EMG signal during REMure in comparison with NREMure (Clement *et al*., 2008). Horner and Kubin (1999) demonstrated that the microinjection of carbachol into the pontine reticular formation potentiates the REM-like state in rats anesthetized with urethane, evoking clear muscle atonia (Horner & Kubin, 1999); however, when those rats were sensory stimulated (pinch in the hindlimb) they showed an EEG with theta oscillations similar to REM sleep but with an increase in muscle tone.

We have demonstrated that urethane anesthesia and physiological sleep differ in the duration of the alternating cycles (Figure 1 E, F and G). In the natural sleep cycle, REM sleep represented 11.6% of the total time, while NREM represented 88.4%. During the anesthetized states, the time spent in NREMure and REMure was almost evenly divided at 45.73% Vs. 54.27%, respectively. This was due to equivalent number and duration of bouts in each state. This is in accordance with other studies that found similar amounts of REMure and NREMure during urethane anesthesia (Pagliardini *et al*., 2012). Interestingly, Clement et al. (2008) found that an increase in the depth of anesthesia reduces the time spent in REMure while the cyclicity is maintained (Clement *et al*., 2008).

### EEG during urethane anesthesia

We determined that urethane anesthesia affects the spectral power of the EEG in different frequency bands. Clear differences were found between W and the anesthetized states. Compared to the waking state, NREMure had a large increase in delta power as well as a reduction in power across all other frequency bands (theta, sigma, beta, LG, HG and HFO). Additionally, W was characterized by a higher power than REMure, especially in the higher frequency bands (HFO in rOB, rM1, rS1; HG and LG in rM1 and rS1 and beta in rS1). This is particularly interesting because higher frequency bands have been associated with consciousness and cognitive processing (Kucewicz *et al*., 2014; Pal *et al*., 2016; Castro-Zaballa *et al*., 2018).

The spectral power during NREMure also differs from physiological NREM sleep. Lower delta was higher in NREMure, while at approximately 3 Hz, NREM sleep had similar or even higher power than NREMure. Since EEG oscillations follow a power-law scaling, by which slower oscillations have higher power values (Miller *et al*., 2009), it is possible that during NREMure there was a shift in the peak frequency of the delta oscillations, with slower delta waves during the anesthetized states in comparison with natural NREM sleep. Interestingly, while delta oscillations during anesthesia with propofol or isoflurane have been shown to have a similar amplitude than NREM sleep delta oscillations (Murphy *et al*., 2011; Akeju & Brown, 2017), the frequency of delta oscillations is slower during anesthesia with ketamine and medetomidine (Torao-Angosto *et al*., 2021).

On the other hand, the power of higher frequency bands was higher in NREM sleep in most cortical locations (see below). Sigma band (10-15 Hz) encompasses the frequency range of sleep spindles, a characteristic electroencephalographic feature of NREM sleep. In fact, NREM sleep is characterized by a higher sigma power than W and REM sleep (Mondino *et al*., 2020). In this study, NREMure showed lower sigma power values than NREM sleep (Figure 2B) and lower than REMure (Figure 2A). Interestingly, Murphy *et* al. (2011) showed in humans that while NREM sleep and propofol anesthesia EEG characteristics shared several similarities, propofol failed to effectively generate spindles (Murphy *et al*., 2011). In the present study the number and duration of sleep spindles was not calculated, but Clement et al. (2008) showed that sleep spindles did occur during urethane anesthesia (Clement *et al*., 2008). However, this study found that spindles occur more frequently during transitions between NREMure and REMure, which is also seen during spontaneous NREM-REM transitions (Bandarabadi *et al*., 2020). These transitional epochs were not considered in the present study.

When comparing REMure with REM sleep, we found that delta power was higher during REMure. It is interesting that in both states of urethane, delta power was higher than during their analogous state in the physiological sleep. Of note, an increase in delta oscillations characterizes general anesthesia with GABA agents (Hagihira, 2015; Mondino *et al*., 2020). On the contrary, the power of most other frequency bands was larger in REM sleep than in REMure, mainly in anterior locations. This could be related to the intense cognitive processing associated with sleep and possibly related to memory consolidation.

We observed a clear decrease of the peak of the theta band frequency under REMure. In fact, Fenik and Kubin (2009) have already described that hippocampal theta-like rhythms had a lower frequency range in rats anesthetized with urethane (3-5 Hz) than in awake rats (6-8 Hz); however, Dringenber and Vanderwolf (1998) and Vertes (1984), suggested that these rhythms are generated by similar neuronal networks and that can be triggered by similar stimuli (Vertes, 1984; Dringenberg & Vanderwolf, 1998; Fenik & Kubin, 2009). Clement et al. (2008) found that the spindles under urethane anesthesia also have a lower frequency (≈8Hz; i.e., in the theta frequency band range) than physiological sleep spindles (Clement *et al*., 2008). However, the mechanisms of the reduction in the theta or spindles frequency are not understood. Our results are consistent with previous studies identifying a decreased frequency of hippocampal theta during isoflurane anesthesia (Perouansky *et al*., 2007; Mashour *et al*., 2010)

Additionally, we observed a decrease in power of the high frequency oscillations in both NREMure and REMure. This finding is typically associated with NREM sleep and general anesthesia with isoflurane, but not with ketamine (a NMDA receptor antagonist) (Kortelainen *et al*., 2012; Akeju *et al*., 2016; Castro-Zaballa *et al*., 2018; Mondino *et al*., 2020). Of note, a clear peak in HFO power, suggesting a true HFO oscillation during REM sleep have been described in the OB and posterior cortices (Cavelli *et al*., 2018; Mondino *et al*., 2020).

We also analyzed the inter and intra-hemispheric coherence. When comparing W and both anesthetized states, delta z’coherence was larger in NREMure, primarily for interhemispheric derivatives, suggesting the presence of synchronized slow wave activity in both hemispheres. By contrast, high frequency (HG and HFO) z’coherence was larger during W reaching significance for S1 interhemispheric, M1-S1 and S1-V2 (only for gamma) derivatives. Interestingly, high frequency z’coherence is lost during both GABA agents and ketamine general anesthesia (Purdon *et al*., 2013; Pal *et al*., 2015), as well as during NREM and REM sleep (Cavelli *et al*., 2015; Cavelli *et al*., 2017; Castro-Zaballa *et al*., 2018).

We also analyzed the anteroposterior (rM1–rV2) directed connectivity by means of NSTE. We demonstrated that under NREMure there was a decrease of theta connectivity both in FF and FB direction. Furthermore, in comparison to W, REMure showed a reduction of HG band connectivity in the FB directionality but not in the FF, which is consistent with the reduction or loss in gamma FB transfer entropy during sleep and anesthesia with different agents (Lee *et al*., 2013; Pal *et al*., 2016; Li *et al*., 2017).

EEG complexity was also analyzed using the Lempel-Ziv algorithm. Compared to W, LZC was reduced in most cortical locations in both anesthetized states; the decrease in complexity was larger during NREMure than REMure, similar to what has been shown to occur with physiological sleep states (Abásolo *et al*., 2015). In addition, when only low frequencies (1-15 Hz) were analyzed, NREMure and REMure had reduced complexity in comparison to their analogous natural states. In this regard, Gonzalez et al. (2019) have shown that complexity, assessed by permutation entropy, is higher during W than both sleep states (Gonzalez *et al*., 2019). Furthermore, EEG complexity decreases under general anesthesia (Zhang *et al*., 2001; Hudetz *et al*., 2016). Notably, in comparison to W, we also found a decrease in complexity during NREMure and REMure assessed by permutation entropy (data not shown).

We also introduced the novel analysis of JLZC, which provides a broader view of cortical complexity. For example, homogenous cortical activity results in low JLZC while the opposite occurs when the activity is differentiated across cortical areas. This analysis showed that, in comparison to W, JLZC decreases during both anesthetized states, but the decrease is larger during NREMure (Mateos *et al*., 2017). Moreover, if we isolated low-frequency activity, JLZC is lower under urethane in comparison to the analogous natural sleep states. Interestingly, Mateos et. al (2017) analyzed standard EEG in humans and found that, at low frequencies (4-12 Hz), JLZC was lower during NREM than during W, but JLZC during REM sleep was similar or even higher than during W. This could be associated with the cognitive processes that takes place during REM sleep (Hobson, 2009). These authors also found that JLZC was exceptionally low during comatose states and epileptic seizures with loss of consciousness, showing that JLZC is an important candidate as a correlate of consciousness. According to these findings, we could assume that the state of consciousness is reduced during urethane anesthesia in comparison with its analog during sleep. To our knowledge, this is the first report of the effect of an anesthetic on JLZC.

### EEG correlates of consciousness are lost in urethane anesthesia

The concept of “neural correlates of consciousness” (NCC) represents the minimal set of neural events and structures sufficient to generate any kind of experience (Koch *et al*., 2016; Mashour & Hudetz, 2018; Torterolo *et al*., 2019; Sarasso *et al*., 2021). In a very schematic way, the main EEG correlates of W consciousness, and the modifications under NREMure and REMure are listed in Table 1. W presents a relative low delta, as well as high gamma and HFO power and coherence in the EEG. While all these signatures are altered by NREMure, under REMure high frequency oscillations power and coherence are lost in most cortical locations and derivates. During REMure there was a decrease in the HG FB directionality of the signal, a feature that has been established as a correlate of loss of consciousness (Pal *et al*., 2016). Interestingly, while the values during NREMure were lower than during W, significance was not achieved. Finally, high EEG complexity also characterizes normal W and is decreased during both NREMure and REMure; high complexity is also considered a marker of consciousness (Abásolo *et al*., 2015; Mateos *et al*., 2017). Hence, EEG features strongly suggest that urethane produces unconsciousness not only under NREMure, that have similarities with the EEG effect of GABA anesthetic agents and deep NREM sleep, but also under the activated state of REMure.

**Table 1.** Parameters associated with consciousness/unconsciousness during anesthetized states and physiological sleep, in comparison to wakefulness.

|  | NREMure | NREM sleep | REMure | REM sleep |
| --- | --- | --- | --- | --- |
| Delta power | ↑ ↑ | ↑<sup>a</sup> | ↑ | n/c<sup>a</sup> |
| Gamma and HFO power | ↓ ↓ | ↓<sup>a</sup> | ↓ | n/c<sup>a</sup> |
| Gamma and HFO coherence | ↓ | ↓<sup>a</sup> | ↓ | ↓<sup>a</sup> |
| HG FB Gamma NSTE | n/c | ↓<sup>b</sup> | ↓ | ↓<sup>b</sup> |
| LZC | ↓ | ↓<sup>c</sup> | ↓ | n/c<sup>c</sup> |
| JLZC | ↓ | ↓<sup>d</sup> | ↓ | n/c<sup>d</sup> |

EEG features suggest that urethane produces unconsciousness during both NREMure and REMure. Increase (+) or decrease (-) in comparison to wakefulness. n/c, no changes. <sup>a</sup>Mondino et. al (2020), <sup>b</sup>Pal et. al (2016), <sup>c</sup>Abásolo et. al (2015), <sup>d</sup>Mateos et al. 2017

### Urethane anesthesia is not a faithful model of sleep

Despite the superficial similarity of the EEG between NREMure and natural NREM sleep, as well as between REMure and REM sleep, a deeper analysis shows strong differences between physiological and pharmacological states.

NREM sleep, especially during deep NREM sleep or N3 where EEG slow waves are fully developed, is considered a state where oneiric behavior is scarce (Siclari *et al*., 2017). However, NREM sleep has a critical function in cognitive functions such as memory consolidation (Stickgold, 2005; Berner *et al*., 2006; Wei *et al*., 2018). The EEG profile of NREMure strongly differs from natural NREM sleep. Compared to NREM sleep, NREMure exhibits a potentiation of delta waves power associated with a decrease in higher frequency power. In addition, NREMure shows a decrease in theta to gamma inter-hemispheric coherence in V2 and/or S1 derivates. Finally, considering only the lower (< 15 Hz) frequency oscillations, we also found that NREMure has lower LZC and JLZC.

REM sleep is an activated state where the neural activity and neurochemical milieu create conditions for the emergence of phenomenal experience in the form of dreams (Hobson, 2009; Siclari *et al*., 2017). The EEG profile under REMure also differs from the natural counterpart. Compared to REM sleep, REMure showed a clear decrease in theta oscillation frequency more consistent with an anesthetized state. Moreover, while delta power was higher in REMure, the power of higher frequencies decreased in this anesthetized state, mainly in anterior cortices. Decreases in interhemispheric coherence in theta and sigma band during REMure were also detected in posterior cortices, while an increase in HFO in S1 interhemispheric coherence was also observed. As in NREMure, a decrease in complexity (both LZC and JLZC) was also detected.

To summarize, these analyses demonstrate that, despite their similarities, the EEG in urethane-anaesthetized states differs from sleep. Hence it is possible that cognitive functions that are known to be carried on during sleep, such as memory consolidation (Stickgold, 2005; Hobson, 2009), are not taking place under urethane anesthesia. Of note, intense research has been carried out trying to uncover the precise mechanisms by which anesthetics induce unconsciousness. In the past two decades there has been a focus on the shared mechanisms between anesthesia and sleep with a body of correlative suggesting that anesthetics and sleep share the same neural circuits. However, in a recent study, we have demonstrated that the activation of GABAergic or glutamatergic neurons in the preoptic area of the hypothalamus, a key area for sleep generation, alters sleep-wake architecture but does not modulate the anesthetized state (Vanini *et al*., 2020). The results from this paper can be considered as additional evidence against a strong mechanistic overlap between sleep and anesthesia. This is especially compelling because urethane was considered to be the only general anesthetic that could mimic the full spectrum of natural sleep. However, despite the similarities between urethane anesthesia and sleep, this detailed analysis revealed key differences.

### Limitations

Our study has several limitations, first, we have not evaluated different doses of urethane anesthesia; therefore, lighter or deeper planes of anesthesia could show different results. Additionally, anesthesia was achieved by a single intraperitoneal injection of urethane and no supplementary injections were given, therefore, the anesthetic effects could have been attenuated over time. However, other authors evaluated the effect of additional injections over time and showed that urethane cyclicity is not affected by the absolute concentration of urethane in blood(Clement *et al*., 2008). Finally, this study was performed with male rats, and therefore we could be losing some sex-differences of urethane anesthesia.

## Conclusions

Urethane produces two different alternating electrographic states: a NREM sleep -like state and an activated state that has features of either REM sleep or waking. Our study revealed EEG signatures of unconsciousness in both urethane-induced states. In addition, NREM-like and REM-like states during urethane anesthesia differ from natural sleep states.

## References

Abásolo, D., Simons, S., Morgado da Silva, R., Tononi, G. & Vyazovskiy, V.V. (2015) Lempel-Ziv complexity of cortical activity during sleep and waking in rats. Journal of neurophysiology, 113, 2742–2752.

Akeju, O. & Brown, E.N. (2017) Neural oscillations demonstrate that general anesthesia and sedative states are neurophysiologically distinct from sleep. Curr Opin Neurobiol, 44, 178–185.

Akeju, O., Song, A.H., Hamilos, A.E., Pavone, K.J., Flores, F.J., Brown, E.N. & Purdon, P.L. (2016) Electroencephalogram signatures of ketamine anesthesia-induced unconsciousness. Clin Neurophysiol, 127, 2414–2422.

Arena, A., Comolatti, R., Thon, S., Casali, A.G. & Storm, J.F. (2021) General Anesthesia Disrupts Complex Cortical Dynamics in Response to Intracranial Electrical Stimulation in Rats. eNeuro, 8.

Bandarabadi, M., Herrera, C.G., Gent, T.C., Bassetti, C., Schindler, K. & Adamantidis, A.R. (2020) A role for spindles in the onset of rapid eye movement sleep. Nat Commun, 11, 5247.

Berner, I., Schabus, M., Wienerroither, T. & Klimesch, W. (2006) The significance of sigma neurofeedback training on sleep spindles and aspects of declarative memory. Appl Psychophysiol Biofeedback, 31, 97–114.

Borjigin, J., Lee, U., Liu, T., Pal, D., Huff, S., Klarr, D., Sloboda, J., Hernandez, J., Wang, M.M. & Mashour, G.A. (2013) Surge of neurophysiological coherence and connectivity in the dying brain. Proc Natl Acad Sci U S A, 110, 14432–14437.

Bullock, T. & McClune, M. (1989) Lateral coherence of the electrocorticogram: a new measure of brain synchrony. Electroencephalography and clinical Neurophysiology, 73, 479–498.

Castro-Zaballa, S., Cavelli, M.L., Gonzalez, J., Nardi, A.E., Machado, S., Scorza, C. & Torterolo, P. (2018) EEG 40 Hz Coherence Decreases in REM Sleep and Ketamine Model of Psychosis. Front Psychiatry, 9, 766.

Cavelli, M., Castro-Zaballa, S., Mondino, A., Gonzalez, J., Falconi, A. & Torterolo, P. (2017) Absence of EEG gamma coherence in a local activated cortical state: a conserved trait of REM sleep. Translational Brain Rhythmicity, 2.

Cavelli, M., Castro, S., Schwarzkopf, N., Chase, M.H., Falconi, A. & Torterolo, P. (2015) Coherent neocortical gamma oscillations decrease during REM sleep in the rat. Behav Brain Res, 281, 318–325.

Cavelli, M., Rojas-Libano, D., Schwarzkopf, N., Castro-Zaballa, S., Gonzalez, J., Mondino, A., Santana, N., Benedetto, L., Falconi, A. & Torterolo, P. (2018) Power and coherence of cortical high-frequency oscillations during wakefulness and sleep. Eur J Neurosci, 48, 2728–2737.

Clement, E.A., Richard, A., Thwaites, M., Ailon, J., Peters, S. & Dickson, C.T. (2008) Cyclic and sleep-like spontaneous alternations of brain state under urethane anaesthesia. PLoS One, 3, e2004.

Demertzi, A., Tagliazucchi, E., Dehaene, S., Deco, G., Barttfeld, P., Raimondo, F., Martial, C., Fernández-Espejo, D., Rohaut, B., Voss, H.U., Schiff, N.D., Owen, A.M., Laureys, S., Naccache, L. & Sitt, J.D. (2019) Human consciousness is supported by dynamic complex patterns of brain signal coordination. 5, eaat7603.

Détári, L. & Vanderwolf, C.H. (1987) Activity of identified cortically projecting and other basal forebrain neurones during large slow waves and cortical activation in anaesthetized rats. Brain Research, 437, 1–8.

Dringenberg, H.C. & Vanderwolf, C.H. (1998) Involvement of Direct and Indirect Pathways in Electrocorticographic Activation. Neuroscience & Biobehavioral Reviews, 22, 243–257.

Fenik, V.B. & Kubin, L. (2009) Differential localization of carbachol-and bicuculline-sensitive pontine sites for eliciting REM sleep-like effects in anesthetized rats. J Sleep Res, 18, 99–112.

González, J., Cavelli, M., Castro-Zaballa, S., Mondino, A., Tort, A.B.L., Rubido, N., Carrera, I. & Torterolo, P. (2021) EEG Gamma Band Alterations and REM-like Traits Underpin the Acute Effect of the Atypical Psychedelic Ibogaine in the Rat. ACS Pharmacology & Translational Science.

Gonzalez, J., Cavelli, M., Mondino, A., Pascovich, C., Castro-Zaballa, S., Rubido, N. & Torterolo, P. (2020) Electrocortical temporal complexity during wakefulness and sleep: an updated account. Sleep Science, 47–50.

Gonzalez, J., Cavelli, M., Mondino, A., Pascovich, C., Castro-Zaballa, S., Torterolo, P. & Rubido, N. (2019) Decreased electrocortical temporal complexity distinguishes sleep from wakefulness. Sci Rep, 9, 18457.

Gonzalez, J., Prieto, J.P., Rodriguez, P., Cavelli, M., Benedetto, L., Mondino, A., Pazos, M., Seoane, G., Carrera, I., Scorza, C. & Torterolo, P. (2018) Ibogaine Acute Administration in Rats Promotes Wakefulness, Long-Lasting REM Sleep Suppression, and a Distinctive Motor Profile. Front Pharmacol, 9, 374.

Hagihira, S. (2015) Changes in the electroencephalogram during anaesthesia and their physiological basis. Br J Anaesth, 115 Suppl 1, i27–i31.

Hay, Y.A. (2021) Brainstem cholinergic modulation of the thalamocortical activity in urethane anesthetized mice. Preprint bioRxiv.

Hobson, J.A. (2009) REM sleep and dreaming: towards a theory of protoconsciousness. Nat Rev Neurosci, 10, 803–813.

Horner, R.L. & Kubin, L. (1999) Pontine Carbachol Elicits Multiple Rapid Eye Movement Sleep-Like Neural Events in Urethane-Anesthetized Rats. Neuroscience, 93, 215–226.

Hudetz, A.G., Liu, X., Pillay, S., Boly, M. & Tononi, G. (2016) Propofol anesthesia reduces Lempel-Ziv complexity of spontaneous brain activity in rats. Neurosci Lett, 628, 132–135.

Hudetz, A.G., Vizuete, J.A. & Pillay, S. (2011) Differential effects of isoflurane on high-frequency and low-frequency gamma oscillations in the cerebral cortex and hippocampus in freely moving rats. Anesthesiology, 114, 588–595.

Koch, C., Massimini, M., Boly, M. & Tononi, G. (2016) Neural correlates of consciousness: progress and problems. Nature Reviews Neuroscience, 17, 307–321.

Kortelainen, J., Jia, X., Seppanen, T. & Thakor, N. (2012) Increased electroencephalographic gamma activity reveals awakening from isoflurane anaesthesia in rats. Br J Anaesth, 109, 782–789.

Kucewicz, M.T., Cimbalnik, J., Matsumoto, J.Y., Brinkmann, B.H., Bower, M.R., Vasoli, V., Sulc, V., Meyer, F., Marsh, W.R., Stead, S.M. & Worrell, G.A. (2014) High frequency oscillations are associated with cognitive processing in human recognition memory. Brain, 137, 2231–2244.

Lee, U., Ku, S., Noh, G., Baek, S., Choi, B. & Mashour, G.A. (2013) Disruption of frontal-parietal communication by ketamine, propofol, and sevoflurane. Anesthesiology, 118, 1264–1275.

Lempel, A. & Ziv, J. (1976) On the Complexity of Finite Sequences. IEEE Transactions on Information Theory, 22, 75–81.

Li, D., Hambrecht-Wiedbusch, V.S. & Mashour, G.A. (2017) Accelerated Recovery of Consciousness after General Anesthesia Is Associated with Increased Functional Brain Connectivity in the High-Gamma Bandwidth. Front Syst Neurosci, 11, 16.

Maggi, C. & Meli, A. (1986) Suitability of urethane anesthesia for physiopharmacological investigations in various systems. Part 1: General considerations. Experientia, 42, 109–210.

Mashour, G.A. & Hudetz, A.G. (2018) Neural Correlates of Unconsciousness in Large-Scale Brain Networks. Trends Neurosci, 41, 150–160.

Mashour, G.A., Lipinski, W.J., Matlen, L.B., Walker, A.J., Turner, A.M., Schoen, W., Lee, U. & Poe, G.R. (2010) Isoflurane anesthesia does not satisfy the homeostatic need for rapid eye movement sleep. Anesth Analg, 110, 1283–1289.

Mateos, D.M., Wennberg, R., Guevara, R. & Perez Velazquez, J.L. (2017) Consciousness as a global property of brain dynamic activity. Physical Review E, 96, 062410.

Melloni, L., Molina, C., Pena, M., Torres, D., Singer, W. & Rodriguez, E. (2007) Synchronization of neural activity across cortical areas correlates with conscious perception. J Neurosci, 27, 2858–2865.

Miller, K.J., Sorensen, L.B., Ojemann, J.G. & den Nijs, M. (2009) Power-law scaling in the brain surface electric potential. PLoS Comput Biol, 5, e1000609.

Mondino, A., Cavelli, M., Gonzalez, J., Osorio, L., Castro-Zaballa, S., Costa, A., Vanini, G. & Torterolo, P. (2020) Power and Coherence in the EEG of the Rat: Impact of Behavioral States, Cortical Area, Lateralization and Light/Dark Phases. Clocks Sleep, 2, 536–556.

Mondino, A., Cavelli, M., Gonzalez, J., Santana, N., Castro-Zaballa, S., Mechoso, B., Bracesco, N., Fernandez, S., Garcia-Carnelli, C., Castro, M.J., Umpierrez, E., Murillo-Rodriguez, E., Torterolo, P. & Falconi, A. (2019) Acute effect of vaporized Cannabis on sleep and electrocortical activity. Pharmacol Biochem Behav, 179, 113–123.

Mondino, A., Hambrecht-Wiedbusch, V., Li, D., York, A.K., Pal, D., Gonzalez, J., Torterolo, P., Mashour, G.A. & Vanini, G. (2021) Glutamatergic neurons in the preoptic hypothalamus promote wakefulness, destabilize NREM sleep, suppress REM sleep, and regulate cortical dynamics. J Neurosci.

Murakami, M., Kashiwadani, H., Kirino, Y. & Mori, K. (2005) State-dependent sensory gating in olfactory cortex. Neuron, 46, 285–296.

Murphy, M., Bruno, M.A., Riedner, B., Boveroux, P., Noirhomme, Q., Landsness, E., Brichant, J., Phillips, C., Massimini, M., Laureys, S., Tononi, G. & Melanie, B. (2011) Propofol anesthesia and sleep: a high-density EEG study. Sleep, 34, 283–291.

Neves, R.M., van Keulen, S., Yang, M., Logothetis, N.K. & Eschenko, O. (2018) Locus coeruleus phasic discharge is essential for stimulus-induced gamma oscillations in the prefrontal cortex. J Neurophysiol, 119, 904–920.

Pagliardini, S., Gosgnach, S. & Dickson, C.T. (2013) Spontaneous sleep-like brain state alternations and breathing characteristics in urethane anesthetized mice. PLoS One, 8, e70411.

Pagliardini, S., Greer, J.J., Funk, G.D. & Dickson, C.T. (2012) State-dependent modulation of breathing in urethane-anesthetized rats. J Neurosci, 32, 11259–11270.

Pal, D., Hambrecht-Wiedbusch, V.S., Silverstein, B.H. & Mashour, G.A. (2015) Electroencephalographic coherence and cortical acetylcholine during ketamine-induced unconsciousness. Br J Anaesth, 114, 979–989.

Pal, D., Silverstein, B.H., Lee, H. & Mashour, G.A. (2016) Neural Correlates of Wakefulness, Sleep, and General Anesthesia: An Experimental Study in Rat. Anesthesiology, 125, 929–942.

Perouansky, M. H. H., Perkins, M. & Pearce, R.A. (2007) Amnesic concentrations of the nonimmobilizer 1,2-dichlorohexafluorocyclobutane (F6, 2N) and isoflurane alter hippocampal theta oscillations in vivo. Anesthesiology, 106, 1168–1176.

Pigorini, A., Sarasso, S., Proserpio, P., Szymanski, C., Arnulfo, G., Casarotto, S., Fecchio, M., Rosanova, M., Mariotti, M., Lo Russo, G., Palva, J.M., Nobili, L. & Massimini, M. (2015) Bistability breaks-off deterministic responses to intracortical stimulation during non-REM sleep. Neuroimage, 15, 105–113.

Purdon, P.L., Pierce, E.T., Mukamel, E.A., Prerau, M.J., Walsh, J.L., Wong, K.F., Salazar-Gomez, A.F., Harrell, P.G., Sampson, A.L., Cimenser, A., Ching, S., Kopell, N.J., Tavares-Stoeckel, C., Habeeb, K., Merhar, R. & Brown, E.N. (2013) Electroencephalogram signatures of loss and recovery of consciousness from propofol. Proc Natl Acad Sci U S A, 110, E1142–1151.

Sarasso, S., Casali, A.G., Casarotto, S., Rosanova, M., Sinigaglia, C. & M., M. (2021) Consciousness and complexity: a consilience of evidence. Neuroscience of Consciousness, 7, 1–24.

Schartner, M., Seth, A., Noirhomme, Q., Boly, M., Bruno, M.A., Laureys, S. & Barrett, A. (2015) Complexity of Multi-Dimensional Spontaneous EEG Decreases during Propofol Induced General Anaesthesia. PLoS One, 10, e0133532.

Schartner, M.M., Pigorini, A., Gibbs, S.A., Arnulfo, G., Sarasso, S., Barnett, L., Nobili, L., Massimini, M., Seth, A.K. & Barrett, A.B. (2017) Global and local complexity of intracranial EEG decreases during NREM sleep. Neurosci Conscious, 2017, niw022.

Siclari, F., Baird, B., Perogamvros, L., Bernardi, G., LaRocque, J.J., Riedner, B., Boly, M., Postle, B.R. & Tononi, G. (2017) The neural correlates of dreaming. Nat Neurosci, 20, 872–878.

Steriade, M., Nunez, A. & Amzica, F. (1993) A novel slow (< 1 Hz) oscillation of neocortical neurons in vivo: depolarizing and hyperpolarizing components. J Neurosci, 13, 3252–3265.

Stickgold, R. (2005) Sleep-dependent memory consolidation. Nature, 437, 1272–1278.

Tononi, G., Boly, M., Massimini, M. & Koch, C. (2016) Integrated information theory: from consciousness to its physical substrate. Nat Rev Neurosci, 17, 450–461.

Torao-Angosto, M., Manasanch, A., Mattia, M. & Sanchez-Vives, M.V. (2021) Up and Down States During Slow Oscillations in Slow-Wave Sleep and Different Levels of Anesthesia. Front Syst Neurosci, 15, 609645.

Torterolo, P., Castro-Zaballa, S., Cavelli, M. & Gonzalez, J. (2019) Arousal and normal conscious cognition Arousal in Neurological and Psychiatric Diseases, pp. 1–24.

Vanini, G., Bassana, M., Mast, M., Mondino, A., Cerda, I., Phyle, M., Chen, V., Colmenero, A.V., Hambrecht-Wiedbusch, V.S. & Mashour, G.A. (2020) Activation of Preoptic GABAergic or Glutamatergic Neurons Modulates Sleep-Wake Architecture, but Not Anesthetic State Transitions. Curr Biol, 30, 779–787 e774.

Varela, F., Lachaux, J., Rodriguez, E. & Martinerie, J. (2001) The BrainWeb: Phase Sinchronization and Large-Scale Integration. Nature Reviews Neurosciences, 2, 229–239.

Vertes, R. (1984) Brainstem Control of The Events of REM Sleep. Progress in Neurobiology, 22, 241 –288.

Wei, Y., Krishnan, G.P., Komarov, M. & Bazhenov, M. (2018) Differential roles of sleep spindles and sleep slow oscillations in memory consolidation. PLoS Comput Biol, 14, e1006322.

Yagishita, H., Nishimura, Y., Noguchi, A., Shikano, Y., Ikegaya, Y. & Sasaki, T. (2020) Urethane anesthesia suppresses hippocampal subthreshold activity and neuronal synchronization. Brain Res, 1749, 147137.

Zhang, X., Roy, R. & Jensen, E. (2001) EEG Complexity as a Measure of Depth of Anesthesia for Patients. IEEE TRANSACTIONS ON BIOMEDICAL ENGINEERING, 48, 1424–1433.

Zozor, S., Ravier, P. & Buttelli, O. (2005) On Lempel–Ziv complexity for multidimensional data analysis. Physica A: Statistical Mechanics and its Applications, 345, 285–302.

