## Supplementary material for "Urethane Anesthesia Exhibits Neurophysiological Correlates of Unconsciousness and is Distinct from Sleep": Supplementary tables Urethane.docx

|  | **OB** | | | **M1** | | | **S1** | | | **V2** | | |
| --- | --- | --- | --- | --- | --- | --- | --- | --- | --- | --- | --- | --- |
| **Frequency** | **df** | **F** | **p** | **df** | **F** | **p** | **df** | **F** | **p** | **df** | **F** | **p** |
| Delta | 2,7 | 19.24 | 0.001* | 2,8 | 45.87 | <0.001* | 2,8 | 54.90 | <0.001* | 2,7 | 23.04 | 0.001* |
| Theta | 2,7 | 13.77 | 0.005* | 2,8 | 12.26 | 0.004* | 2,8 | 18.19 | 0.002* | 2,7 | 9.28 | 0.009* |
| Sigma | 2,7 | 3.717 | 0.053 | 2,8 | 21.00 | <0.001* | 2,8 | 56.86 | <0.001* | 2,7 | 19.10 | 0.001* |
| Beta | 2,7 | 19.12 | 0.001* | 2,8 | 45.45 | <0.001* | 2,8 | 57.61 | <0.001* | 2,7 | 19.64 | 0.001* |
| LG | 2,7 | 31.49 | 0.001* | 2,8 | 90.21 | <0.001* | 2,8 | 66.21 | <0.001* | 2,7 | 12.25 | 0.008* |
| HG | 2,7 | 24.63 | 0.001* | 2,8 | 115.2 | <0.001* | 2,8 | 74.13 | <0.001* | 2,7 | 26.20 | 0.001* |
| HFO | 2,7 | 29.84 | 0.001* | 2,8 | 45.34 | <0.001* | 2,8 | 30.43 | 0.001* | 2,7 | 20.43 | 0.002* |

**Table 1.** Normalized Power. Repeated measures one-way ANOVA.

Results of the Repeated measures one-way ANOVA comparing W with the anesthetized states (NREMure and REMure). Data are plotted in Figure 2.

|  | **OB** | | | **M1** | | | **S1** | | | **V2** | | |
| --- | --- | --- | --- | --- | --- | --- | --- | --- | --- | --- | --- | --- |
| **Frequency** | **W**  **vs. NREMure** | **W**  **vs. REMure** | **NREMure vs. REMure** | **W**  **vs. NREMure** | **W**  **vs. REMure** | **NREMure vs. REMure** | **W**  **vs. NREMure** | **W**  **vs. REMure** | **NREMure vs. REMure** | **W**  **vs. NREMure** | **W**  **vs. REMure** | **NREMure vs. REMure** |
| Delta | 0.0070* | 0.9570 | 0.0084* | 0.0021* | 0.0791 | 0.0048* | 0.0021* | 0.0905 | 0.0021* | 0.0021* | 0.9288 | 0.0048* |
| Theta | 0.0198* | 0.1191 | 0.0198* | 0.0021* | 0.4682 | 0.0228* | 0.0021* | 0.0905 | 0.0088* | 0.0107* | 0.9288 | 0.0706 |
| Sigma | N/A | N/A | N/A | 0.0027* | 0.0220* | 0.1244 | 0.0021* | 0.0030* | 0.0287* | 0.0040* | 0.1934 | 0.0706 |
| Beta | 0.0243* | 0.3870 | 0.0088* | 0.0021* | 0.1244 | 0.0021* | 0.0021* | 0.0448* | 0.0024* | 0.0040* | 0.9288 | 0.0060* |
| LG | 0.0064* | 0.5882 | 0.0018* | 0.0021* | 0.0098* | 0.0021* | 0.0021* | 0.0287* | 0.0024* | 0.0040* | 0.6448 | 0.0229* |
| HG | 0.0064* | 0.0548 | 0.0109* | 0.0021* | 0.0021* | 0.0022* | 0.0021* | 0.0021* | 0.0021* | 0.0040* | 0.1494 | 0.0104* |
| HFO | 0.0078* | 0.0143* | 0.0034* | 0.0021* | 0.0022* | 0.0021* | 0.0030* | 0.0088* | 0.0088* | 0.0098* | 0.0706 | 0.0706 |

**Table 2.** Normalized Power. Holm-Šídák *post-hoc* test.

Adjusted p values from Holm-Šídák multiple comparison test. We analyzed only frequency bands for which ANOVA was significant. The ones not analyzed are represented by “N/A”. Data are plotted in Figure 2A.

| A. **Normalized power adjusted p values NREMure Vs. NREM** | | | | | | | | | | | | |
| --- | --- | --- | --- | --- | --- | --- | --- | --- | --- | --- | --- | --- |
|  | **rOB** | | | **rM1** | | | **rS1** | | | **rV2** | | |
| **Frequency** | **df** | **t** | **p** | **df** | **t** | **p** | **df** | **t** | **p** | **df** | **t** | **p** |
| Delta | 8 | 8.566 | <0.001* | 8 | 7.273 | <0.001* | 8 | 6.491 | 0.001* | 7 | 6.341 | 0.002* |
| Theta | 8 | 7.602 | <0.001* | 8 | 9.500 | <0.001* | 8 | 6.983 | 0.001* | 7 | 4.789 | 0.007* |
| Sigma | 8 | 8.737 | <0.001* | 8 | 5.669 | 0.002* | 8 | 5.895 | 0.002* | 7 | 7.230 | 0.001* |
| Beta | 8 | 4.604 | 0.010* | 8 | 4.281 | 0.011* | 8 | 4.686 | 0.006* | 7 | 5.430 | 0.005* |
| LG | 8 | 2.980 | 0.041* | 8 | 3.287 | 0.033* | 8 | 4.574 | 0.006* | 7 | 1.503 | 0.007* |
| HG | 8 | 3.622 | 0.025* | 8 | 3.197 | 0.033* | 8 | 3.806 | 0.010* | 7 | 0.403 | 0.8977 |
| HFO | 8 | 2.909 | 0.041* | 8 | 3.169 | 0.033* | 8 | 3.340 | 0.010* | 7 | 0.138 | 0.8977 |
| B. **Normalized power adjusted p values REMure Vs. REM** | | | | | | | | | | | | |
|  | **rOB** | | | **rM1** | | | **rS1** | | | **rV2** | | |
| **Frequency** | **df** | **t** | **p** | **df** | **t** | **P** | **df** | **t** | **p** | **df** | **t** | **p** |
| Delta | 8 | 3.722 | 0.029* | 8 | 6.110 | 0.001* | 8 | 5.581 | 0.002* | 7 | 4.621 | 0.014* |
| Theta | 8 | 0.688 | 0.513 | 8 | 1.171 | 0.475 | 8 | 3.404 | 0.018* | 7 | 4.349 | 0.017* |
| Sigma | 8 | 4.333 | 0.017* | 8 | 7.396 | 0.001* | 8 | 8.399 | 0.001* | 7 | 10.12 | 0.001* |
| Beta | 8 | 5.900 | 0.004* | 8 | 13.63 | 0.001* | 8 | 5.038 | 0.005* | 7 | 0.235 | 0.958 |
| LG | 8 | 6.415 | 0.003* | 8 | 6.380 | 0.001* | 8 | 4.255 | 0.008* | 7 | 0.440 | 0.824 |
| HG | 8 | 2.808 | 0.077 | 8 | 5.528 | 0.002* | 8 | 4.831 | 0.005* | 7 | 0.794 | 0.958 |
| HFO | 8 | 2.681 | 0.077 | 8 | 0.923 | 0.475 | 8 | 1.361 | 0.210* | 7 | 0.307 | 0.770 |

**Table 3.** Normalized Power. Comparison between anesthetized states and NREM and REM sleep

Statistical results from two-tailed paired Student t tests comparing NREMure with NREM sleep (A) and REMure with REM sleep (B). p values were adjusted by Holm-Šídák multiple comparison test. Data are plotted in Figure 2B and 2C respectively.

**Table 4.** Inter-hemispheric and Intra-hemispheric z’coherence.

| A. **Inter-hemispheric z’coherence** | | | | | | | | | |
| --- | --- | --- | --- | --- | --- | --- | --- | --- | --- |
|  | **rM1-lM1** | | | **rS1-lS1** | | | **rV2-lV2** | | |
| **Frequency** | **df** | **F** | **p** | **df** | **F** | **P** | **df** | **F** | **p** |
| Delta | 2,7 | 14.66 | 0.001* | 2,7 | 15.80 | 0.001* | 2,7 | 2.876 | 0.113 |
| Theta | 2,7 | 2.711 | 0.131 | 2,7 | 9.36 | 0.007* | 2,7 | 6.458 | 0.027* |
| Sigma | 2,7 | 2.515 | 0.152 | 2,7 | 26.16 | 0.001* | 2,7 | 5.624 | 0.042* |
| Beta | 2,7 | 0.353 | 0.588 | 2,7 | 8.086 | 0.005* | 2,7 | 6.524 | 0.023* |
| LG | 2,7 | 1.436 | 0.270 | 2,7 | 1.179 | 0.333 | 2,7 | 4.058 | 0.048* |
| HG | 2,7 | 2.635 | 0.141 | 2,7 | 37.30 | <0.001* | 2,7 | 10.24 | 0.012* |
| HFO | 2,7 | 0.311 | 0.672 | 2,7 | 32.77 | <0.001* | 2,7 | 3.789 | 0.088 |
| B. **Intra-hemispheric z’coherence** | | | | | | | | | |
|  | **rOB-rM1** | | | **rM1-rS1** | | | **rS1-rV2** | | |
| **Frequency** | **df** | **F** | **p** | **df** | **F** | **P** | **df** | **F** | **P** |
| Delta | 2,7 | 11.77 | 0.007* | 2,8 | 3.758 | 0.067 | 2,6 | 3.040 | 0.123 |
| Theta | 2,7 | 7.756 | 0.007* | 2,8 | 0.434 | 0.564 | 2,6 | 4.956 | 0.046* |
| Sigma | 2,7 | 7.300 | 0.007* | 2,8 | 1.206 | 0.308 | 2,6 | 0.813 | 0.407 |
| Beta | 2,7 | 3.841 | 0.071 | 2,8 | 0.085 | 0.794 | 2,6 | 0.295 | 0.639 |
| LG | 2,7 | 1.201 | 0.326 | 2,8 | 1.034 | 0.345 | 2,6 | 0.286 | 0.684 |
| HG | 2,7 | 1.158 | 0.325 | 2,8 | 23.04 | <0.001* | 2,6 | 12.29 | 0.004* |
| HFO | 2,7 | 3.217 | 0.094 | 2,8 | 85.11 | <0.001* | 2,6 | 7.393 | 0.012* |

Inter-hemispheric and Intra-hemispheric z’coherence. Comparison between wakefulness and anesthetized states. Repeated measures one-way ANOVA. Data are plotted in Figure 3 and 4.

|  | **rM1-lM1** | | | **rS1-lS1** | | | **rV2-lV2** | | |
| --- | --- | --- | --- | --- | --- | --- | --- | --- | --- |
| **Frequency** | **W**  **vs. NREMure** | **W**  **vs. REMure** | **NREMure vs. REMure** | **W**  **vs. NREMure** | **W**  **vs. REMure** | **NREMure vs. REMure** | **W**  **vs. NREMure** | **W**  **vs. REMure** | **NREMure vs. REMure** |
| Delta | 0.018* | 0.732 | 0.050* | 0.002* | 0.258 | 0.205 | N/A | N/A | N/A |
| Theta | N/A | N/A | N/A | 0.087 | 0.026* | 0.772 | 0.151 | 0.399 | 0.814 |
| Sigma | N/A | N/A | N/A | 0.064 | 0.002* | 0.288 | 0.290 | 0.400 | 0.814 |
| Beta | N/A | N/A | N/A | 0.104 | 0.104 | 0.928 | 0.399 | 0.241 | 0.725 |
| LG | N/A | N/A | N/A | N/A | N/A | N/A | 0.399 | 0.399 | 0.814 |
| HG | N/A | N/A | N/A | 0.002* | 0.007* | 0.894 | 0.131 | 0.233 | 0.814 |
| HFO | N/A | N/A | N/A | 0.003* | 0.007* | 0.737 | N/A | N/A | N/A |

**Table 5.** Inter-hemispheric z’coherence between wakefulness and anesthetized states. Holm-Šídák *post-hoc*.

Adjusted p values from Holm-Šídák multiple comparison test. We analyzed only frequency bands for which ANOVA was significant. The ones not analyzed are represented by “N/A”. Data are plotted in Figure 3.

|  | **rOB-rM1** | | | **rM1-rS1** | | | **rS1-rV2** | | |
| --- | --- | --- | --- | --- | --- | --- | --- | --- | --- |
| **Frequency** | **W**  **vs. NREMure** | **W**  **vs. REMure** | **NREMure vs. REMure** | **W**  **vs. NREMure** | **W**  **vs. REMure** | **NREMure vs. REMure** | **W**  **vs. NREMure** | **W**  **vs. REMure** | **NREMure vs. REMure** |
| Delta | 0.045* | 0.223 | 0.008* | N/A | N/A | 0.205 | N/A | N/A | N/A |
| Theta | 0.065 | 0.223 | 0.223 | N/A | N/A | 0.772 | 0.154 | 0.845 | 0.077 |
| Sigma | 0.106 | 0.785 | 0.065 | N/A | N/A | 0.288 | N/A | N/A | N/A |
| Beta | N/A | N/A | N/A | N/A | N/A | 0.928 | N/A | N/A | N/A |
| LG | N/A | N/A | N/A | N/A | N/A | N/A | N/A | N/A | N/A |
| HG | N/A | N/A | N/A | 0.001* | 0.004* | 0.045* | 0.077 | 0.233 | 0.999 |
| HFO | N/A | N/A | N/A | 0.001* | 0.004* | 0.017* | 0.153 | 0.064 | 0.996 |

**Table 6**. Intra-hemispheric z’coherence. between wakefulness and anesthetized states. Holm-Šídák post-hoc.

Adjusted p values from Holm-Šídák multiple comparison test. We analyzed only frequency bands for which ANOVA was significant. The ones not analyzed are represented by “N/A”. Data are plotted in Figure 4.

**Table 7.** Inter-hemispheric z’coherence between NREMure and NREM, and REMure and REM

| A. **Inter-hemispheric z’coherence p values NREMure Vs. NREM** | | | | | | | | | |
| --- | --- | --- | --- | --- | --- | --- | --- | --- | --- |
|  | **rM1-lM1** | | | **rS1-lS1** | | | **rV2-lV2** | | |
| **Frequency** | **df** | **t** | **P** | **df** | **t** | **P** | **df** | **t** | **p** |
| Delta | 7 | 0.952 | 0.915 | 8 | 0.913 | 0.406 | 7 | 0.521 | 0.854 |
| Theta | 7 | 0.114 | 0.991 | 8 | 3.795 | 0.026 | 7 | 4.988 | 0.011 |
| Sigma | 7 | 0.123 | 0.991 | 8 | 7.540 | 0.001 | 7 | 4.675 | 0.014 |
| Beta | 7 | 1.022 | 0.916 | 8 | 5.417 | 0.005 | 7 | 3.714 | 0.037 |
| LG | 7 | 0.958 | 0.916 | 8 | 6.142 | 0.003 | 7 | 2.875 | 0.092 |
| HG | 7 | 1.029 | 0.916 | 8 | 3.806 | 0.026 | 7 | 0.844 | 0.811 |
| HFO | 7 | 1.461 | 0.766 | 8 | 1.317 | 0.406 | 7 | 0.373 | 0.854 |
| B. **Inter-hemispheric z’coherence p values REMure Vs. REM** | | | | | | | | | |
|  | **rM1-lM1** | | | **rS1-lS1** | | | **rV2-lV2** | | |
| **Frequency** | **df** | **t** | **p** | **df** | **t** | **P** | **df** | **t** | **p** |
| Delta | 7 | 0.120 | 0.907 | 8 | 2.435 | 0.169 | 7 | 0.127 | 0976 |
| Theta | 7 | 0.969 | 0.597 | 8 | 6.023 | 0.003 | 7 | 4.403 | 0.018 |
| Sigma | 7 | 2.309 | 0.323 | 8 | 7.148 | 0.001 | 7 | 5.203 | 0.008 |
| Beta | 7 | 0.157 | 0.534 | 8 | 0.990 | 0.584 | 7 | 3.089 | 0.085 |
| LG | 7 | 0.227 | 0.538 | 8 | 0.060 | 0.953 | 7 | 1.160 | 0.634 |
| HG | 7 | 0.124 | 0.534 | 8 | 1.430 | 0.480 | 7 | 0.204 | 0.976 |
| HFO | 7 | 0.119 | 0.534 | 8 | 3.699 | 0.038 | 7 | 2.000 | 0.301 |

Statistical results from two-tailed paired Student t tests between NREMure and NREM (A) and REMure and REM(B). Adjusted p values (Holm-Šídák correction for multiple comparisons). Data are plotted in Figure 3.

**Table 8**. Intra-hemispheric z’coherence between NREMure and NREM, and REMure and REM.

| A. **Intra-hemispheric z’coherence p values NREMure Vs. NREM** | | | | | | | | | |
| --- | --- | --- | --- | --- | --- | --- | --- | --- | --- |
|  | **rOB-rM1** | | | **rM1-rS1** | | | **rS1-rV2** | | |
| **Frequency** | **df** | **T** | **p** | **df** | **T** | **p** | **df** | **T** | **p** |
| Delta | 7 | 0.876 | 0.558 | 8 | 0.199 | 0.976 | 6 | 0.231 | 0.985 |
| Theta | 7 | 1.288 | 0.558 | 8 | 1.150 | 0.820 | 6 | 0.326 | 0.985 |
| Sigma | 7 | 1.053 | 0.558 | 8 | 1.037 | 0.820 | 6 | 1.331 | 0.842 |
| Beta | 7 | 2.057 | 0.437 | 8 | 1.144 | 0.820 | 6 | 0.726 | 0.942 |
| LG | 7 | 2.012 | 0.437 | 8 | 0.133 | 0.976 | 6 | 0.836 | 0.942 |
| HG | 7 | 1.608 | 0.483 | 8 | 1.243 | 0.820 | 6 | 0.297 | 0.985 |
| HFO | 7 | 1.769 | 0.473 | 8 | 1.557 | 0.700 | 6 | 1.128 | 0.885 |
| B. **Intra-hemispheric z’coherence p values REMure Vs. REM** | | | | | | | | | |
|  | **rOB-rM1** | | | **rM1-rS1** | | | **rS1-rV2** | | |
| **Frequency** | **df** | **T** | **p** | **df** | **T** | **P** | **df** | T | **p** |
| Delta | 7 | 2.802 | 0.149 | 8 | 1.072 | 0.911 | 6 | 3.141 | 0.132 |
| Theta | 7 | 1.475 | 0.456 | 8 | 0.594 | 0.985 | 6 | 2.282 | 0.321 |
| Sigma | 7 | 3.473 | 0.071 | 8 | 0.402 | 0.992 | 6 | 2.035 | 0.369 |
| Beta | 7 | 1.986 | 0.364 | 8 | 0.204 | 0.992 | 6 | 0.371 | 0.979 |
| LG | 7 | 0.255 | 0.806 | 8 | 0.376 | 0.992 | 6 | 0.063 | 0.998 |
| HG | 7 | 1.124 | 0.507 | 8 | 0.110 | 0.992 | 6 | 0.021 | 0.998 |
| HFO | 7 | 1.993 | 0.364 | 8 | 1.129 | 0.911 | 6 | 1.394 | 0.616 |

Statistical results from two-tailed paired Student t tests between NREMure and NREM (A) and REMure and REM(B). Adjusted p values (Holm-Šídák correction for multiple comparisons). Data are plotted in Figure 4.

**Table 9**. Normalized transfer entropy between NREMure and NREM, and REMure and REM.

| **Normalized transfer entropy** | | | | | | |
| --- | --- | --- | --- | --- | --- | --- |
| A. **NREMure vs NREM** | | | | | | |
|  | **FB** | | | **FF** | | |
| **Frequency** | **df** | **t** | **p** | **df** | **t** | **p** |
| Delta | 8 | 0.298 | 0.997 | 8 | 0.572 | 0.970 |
| Theta | 8 | 1.806 | 0.552 | 8 | 1.394 | 0.739 |
| Sigma | 8 | 0.606 | 0.984 | 8 | 1.200 | 0.785 |
| Beta | 8 | 0.127 | 0.997 | 8 | 0.471 | 0.970 |
| LG | 8 | 0.140 | 0.997 | 8 | 0.074 | 0.970 |
| HG | 8 | 0.205 | 0.997 | 8 | 0.571 | 0.970 |
| HFO | 8 | 0.994 | 0.924 | 8 | 1.613 | 0.667 |
| B. **REMure vs REM** | | | | | | |
|  | **FB** | | | **FF** | | |
| **Frequency** | **df** | **t** | **p** | **df** | **t** | **p** |
| Delta | 8 | 0.371 | 0.971 | 8 | 0.406 | 0.9863 |
| Theta | 8 | 0.923 | 0.966 | 8 | 0.435 | 0.9863 |
| Sigma | 8 | 0.923 | 0.966 | 8 | 1.884 | 0.5078 |
| Beta | 8 | 0.962 | 0.971 | 8 | 0.181 | 0.9863 |
| LG | 8 | 0.815 | 0.971 | 8 | 0.075 | 0.9863 |
| HG | 8 | 0.526 | 0.966 | 8 | 1.218 | 0.7748 |
| HFO | 8 | 0.403 | 0.966 | 8 | 1.566 | 0.6385 |

Statistical results from two-tailed paired Student t tests between NREMure and NREM (A) and REMure and REM(B). Adjusted p values (Holm-Šídák correction for multiple comparisons).
